# Tractography from Serial Optical Coherence Tomography: How and Why?

**DOI:** 10.64898/2026.08.14.744847

**Authors:** Charles Poirier, Laurent Petit, Joël Lefebvre, Maxime Descoteaux

**Affiliations:** Department of computer sciences, Université de Sherbrooke, 2500, boul. de l’Université, Sherbrooke, J1K2R1, Qc, Canada; Université Bordeaux, CNRS, CEA, IMN, GIN, UMR 5293, F-33000, Bordeaux, France; Département d’informatique, Université du Québec à Montréal, 201, avenue du Président-Kennedy, Montréal, H2X 3Y7, QC, Canada; IRP/LIA OpTeam, CNRS Biologie et Univ. Bordeaux, France - Univ. Sherbrooke, Sherbrooke, Canada

**Author notes:** Corresponding author (C. Poirier). Co-last authors (Joël Lefebvre, Maxime Descoteaux).

**Keywords:** Serial optical coherence tomography, Microscopy, Orientation distribution function, Tractography

## Abstract

To disentangle complex fiber configurations that remain challenging for diffusion MRI tractography, insights might be gained from microscopy tractography. By precisely following small white matter (WM) fascicles, invisible at the resolution of diffusion MRI, microscopy tractography can help explain how fiber populations are organized at the finest scales. Due to its high resolution and its 3D nature, serial optical coherence tomography (S-OCT) offers promise for studying WM at the microscale. However, whether the reflectivity contrast from S-OCT supports tractography at the microscale is unknown. Furthermore, there is a gap in the literature regarding how an ideal microscopy tractography should behave with respect to the choice of tractography algorithm, tracking maps and microscale orientation distribution functions (µODF) estimation. In this work, we describe a tailored approach for microscopy tractography from S-OCT acquisitions. We validate our approach on a simulated microscopy-like FiberCup dataset, and show that multiscale Frangi filters outperforms structure tensor analysis for estimating µODF. We also show that anatomically-constrained particle filtering tractography enables targeted, region-to-region tractography, and outperforms standard deterministic or probabilistic tracking approaches. We further demonstrate our method on an *ex vivo* whole mouse brain S-OCT reconstruction at 10 µm by reconstructing thalamocortical WM projections. Overall, our results show that SOCT tractography recovers fine WM fascicles that are supported by viral tracing experiments from the Allen Mouse Brain Connectivity Atlas. Moreover, this work shows the first µODF estimation and fully-3D probabilistic particle filtering tractography of the mouse brain from SOCT reconstructions at 10 µm isotropic resolution.

## 1. Introduction

Over the past two decades, many studies have highlighted the need for better anatomical priors for diffusion magnetic resonance imaging (dMRI) fiber tractography (Maier-Hein et al., 2017; Jbabdi and Johansen-Berg, 2011; Girard et al., 2014; Smith et al., 2012). In particular, the inclusion of priors from high-resolution microscopy acquisition is promising for resolving challenging fiber configurations such as fiber crossings and bottlenecks (Descoteaux et al., 2025). Integrating microscopy and tractography has been mostly restricted to validating diffusion MRI metrics (Budde and Frank, 2012; Schilling et al., 2018; Lefebvre et al., 2018) and tractography algorithms (Aydogan et al., 2018; Girard et al., 2020), with some authors using microscopy to improve the reconstruction of fiber orientation distribution functions (FODF) (Liang et al., 2023; Zhu et al., 2025). Following precisely the long-range trajectory of WM fascicles at the microscopic scale is an important step towards the successful integration of microscopy priors into tractography algorithms and downstream connectivity analyses. As such, there has been attempts at performing tractography directly from microscopy acquisitions (Axer et al., 2011; Kjer et al., 2024; Wang et al., 2015; Goergen et al., 2012; Zhu et al., 2025; Liu et al., 2025; Zhang et al., 2025). A few notable approaches are presented below.

In Oh et al. (2014), the authors describe the Allen Mouse Brain Connectivity Atlas (AMBCA), an open-resource for thousands of projection density maps of mouse brain axonal projections acquired in serial 2PE fluorescence microscopy. To obtain a measurable contrast, 2PE fluorescence microsopy requires the injection of neuron-specific virus for labelling neurons located along a region of interest. As such, the measured signal depends on what tracer is used and where it is injected. Also, viral tracers must be allowed time to migrate along the entire brain so they can be measured by the system, increasing sample preparation time. Furthermore, 2PE fluorescence microscopy is limited to 2D images, resulting in highly anisotropic whole brain volumes. Due to its anisotropic nature and lack of orientation information, 2PE fluorescence microscopy viral tracing data from the AMBCA is not suitable for conventional tractography applications. Instead, to connect brain regions, the authors solve for a path maximizing the viral tracer projection density through the voxel grid. Recently, Zhang et al. (2025) showed that high-speed fluorescence volumetric imaging enables the acquisition of Nissl-stained, optically-cleared whole-brain samples at nearly-isotropic micrometer resolution. In their work, the authors show that the organization and shape of Nissl-stained cells can be analyzed to infer the orientation of white matter fascicles, which can then be used for fiber tractography. A downside of this technique is that, like for 2PE fluorescence microscopy, Nissl-staining and optical clearing also increase the sample preparation times. In Sorelli et al. (2023, 2025), the authors apply light-sheet fluorescence microscopy (LSFM) to human brain samples to visualize the myelinated fiber network. At the difference of the previous techniques, LSFM enables fully-3D imaging of samples. It however relies on destructive, time-consuming tissue clearing and staining. To reconstruct fiber orientations from the measured signal, the authors use a Hessian-based image processing pipeline. More recently, tensor light-sheet scattering microscopy (tLSSM) has been suggested to reconstruct orientation maps directly from the system measurements in 3D and in a label-free fashion (Corral-Bolaños et al., 2026). While the method still requires extensive sample preparation times due to optical clearing of brain tissues, the authors show that it enables fiber tractography. The results are however limited to track-density images reconstructed from fiber tractography and lack the trajectory information of individual fibers. Although it is limited to a 2D imaging plane, 3D polarized-light imaging (3D-PLI) also enables brain imaging at the microscale (Axer et al., 2011; Howard et al., 2023; Zhu et al., 2025). 3D-PLI measures the 3D orientation of myelinated fibers without any constrast agents (label-free) or optical clearing. In Zhu et al. (2025), the authors use coregistered PLI and diffusion MRI data from the BigMac dataset (Howard et al., 2023) to estimate hybrid, super-resolved dMRI-PLI FODF for tractography. PLI-only fiber tractography has also been demonstrated inside small samples from a postmortem human brain by locally following the measured local orientation vector (Axer et al., 2011). Finally, polarization-sensitive OCT (PS-OCT) measures the in-plane orientation of myelinated fibers in 3D and without contrast agents (Wang et al., 2015; Liu et al., 2025). As such, PS-OCT has been used for fiber tractography. However, measuring the 3D orientation of fibers requires a complex imaging setup with two imaging objectives. Furthermore, measuring the local orientation from PS-OCT requires integrating the signal along each A-line, resulting in a decrease in the through-plane (axial) resolution.

Conventional serial optical coherence tomography (Lefebvre et al., 2017) (S-OCT) is an intrinsic contrast imaging modality using light interference to measure the reflectivity of a sample. When applied to brain tissues, the measured S-OCT signal is primarily driven by the reflectivity of myelin (Leahy et al., 2013). Its high resolution, in the order of microns, and 3D nature makes it a promising modality for tracking WM fascicles at the microscale. Compared to PS-OCT, conventional S-OCT is less expensive and achieves a higher axial resolution. However, S-OCT does not measure the orientation of WM fibers. A key property of reflectivity-based S-OCT is its orientation-dependent contrast (Lefebvre et al., 2024): fibers parallel to the block face (in-plane) are more visible than fibers oriented along the imaging axis. Consequently, the acquired intensity does not provide a comprehensive view of the underlying WM organization. This is demonstrated in Figure 1 for two mouse brains, where commissural fibers from the corpus callosum (CC) are bright for a coronal acquisition (Figure 1a, white box), and dark for a sagittal acquisition (Figure 1b). While it may seem like a limitation, this orientation-dependency allows for disentangling crossing fiber configurations. For instance, the sagittal acquisition from Figure 1b allows for visualizing WM fibers projecting across the CC from the internal capsule to the cortex (white box). Whether the reflectivity contrast from conventional S-OCT can support the reconstruction of long-range WM fascicles at the microscale remains unknown, with existing examples being limited to *en-face* tracking from small regions of interest (Goergen et al., 2012). In particular, precisely reconstructing microscale WM fascicles with S-OCT tractography requires the careful selection of (a.) an ODF estimation method, and (b.) the tractography algorithm itself.

**Figure 1.**
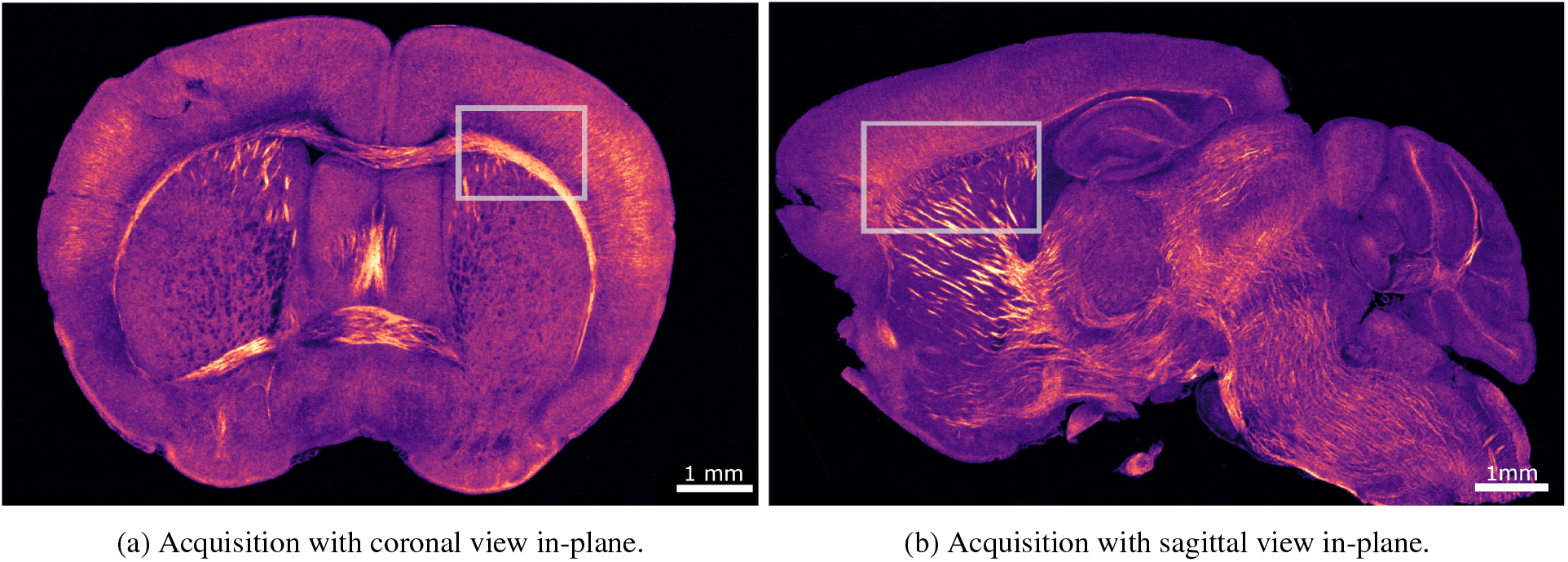
The measured S-OCT signal depends on the orientation of the structure with respect to the imaging objective focal plane. While commissural fibers from the CC come out as very bright when acquired along the coronal plane (a), they do not show up for an acquisition along the sagittal plane (b). The images are shown at a downsampled resolution of 10 µm/px.

*(a*.*) ODF estimation* S-OCT tractography requires a microscopy orientation distribution function (µODF) for driving streamlines reconstruction. When directional information is not directly measured, µODF are estimated by computing a neighbourhood histogram of the local orientations obtained from gradient-based texture analysis (Alberini et al., 2024; Budde and Frank, 2012; Gan and Fleming, 2013; Khan et al., 2015; Kjer et al., 2024; Schilling et al., 2016; Sorelli et al., 2023; Wang et al., 2015; Zhu et al., 2025; Zhang et al., 2025). Recently, Sorelli et al. (2023) described a multiscale approach for estimating the local orientations of myelinated fibers in LSFM, but this approach is not used for fiber tractography. Given that WM fibers vary greatly in diameters, it would be important to study the effect of a multiscale µODF reconstruction method for S-OCT tractography.

*(b*.*) Tractography algorithm* In diffusion MRI, probabilistic tractography approaches allow for better capturing the full extent of hard-to-track WM trajectories by exploring locally less plausible pathways (Jeurissen et al., 2019). While common usage in diffusion MRI tractography, probabilistic approaches are rarely used for microscopy tractography (apart from (Zhu et al., 2025; Zhang et al., 2025)), the assumption being that microscopy voxels are free of partial volume effects: this is however not always the case, with axonal diameters ranging between 0.1 – 10 µm in the human brain (Perge et al., 2012). Also, the inclusion of anatomically-derived rules for dMRI tractography has been shown to greatly reduce the number of false positives by enforcing that the reconstructed streamlines respect some anatomical constraints (Girard et al., 2014; Smith et al., 2012; Aydogan et al., 2018). These anatomical criteria have not yet been implemented for microscopy tractography. However, they could prove useful for accurately reconstructing the trajectories of WM fascicles visible at the microscale.

Here, we develop a tailored methodology for tractography at the microscale from whole mouse brain S-OCT acquisitions. Our approach uses the multiscale Frangi filters from Sorelli et al. (2025) to estimate both a native resolution µODF and a fiber probability map, which are then used as inputs to an anatomically-constrained particle-filtering tractography (Girard et al., 2014) algorithm. The proposed methodology is evaluated on a simulated microscopy-like version of the FiberCup phantom (Fillard et al., 2011; Neher et al., 2017; Côté et al., 2013) and applied to the reconstruction of thalamocortical projections in a whole mouse brain S-OCT volume at 10 µm isotropic resolution. Our results show that, when the imaging plane is properly aligned with respect to the structure of interest, S-OCT tractography enables the reconstruction of WM fascicles at the microscale.

## 2 Methods

This section outlines the steps for performing tractography from a whole-mouse brain S-OCT acquisition. An overview of the proposed pipeline is given in Figure 2.

**Figure 2.**
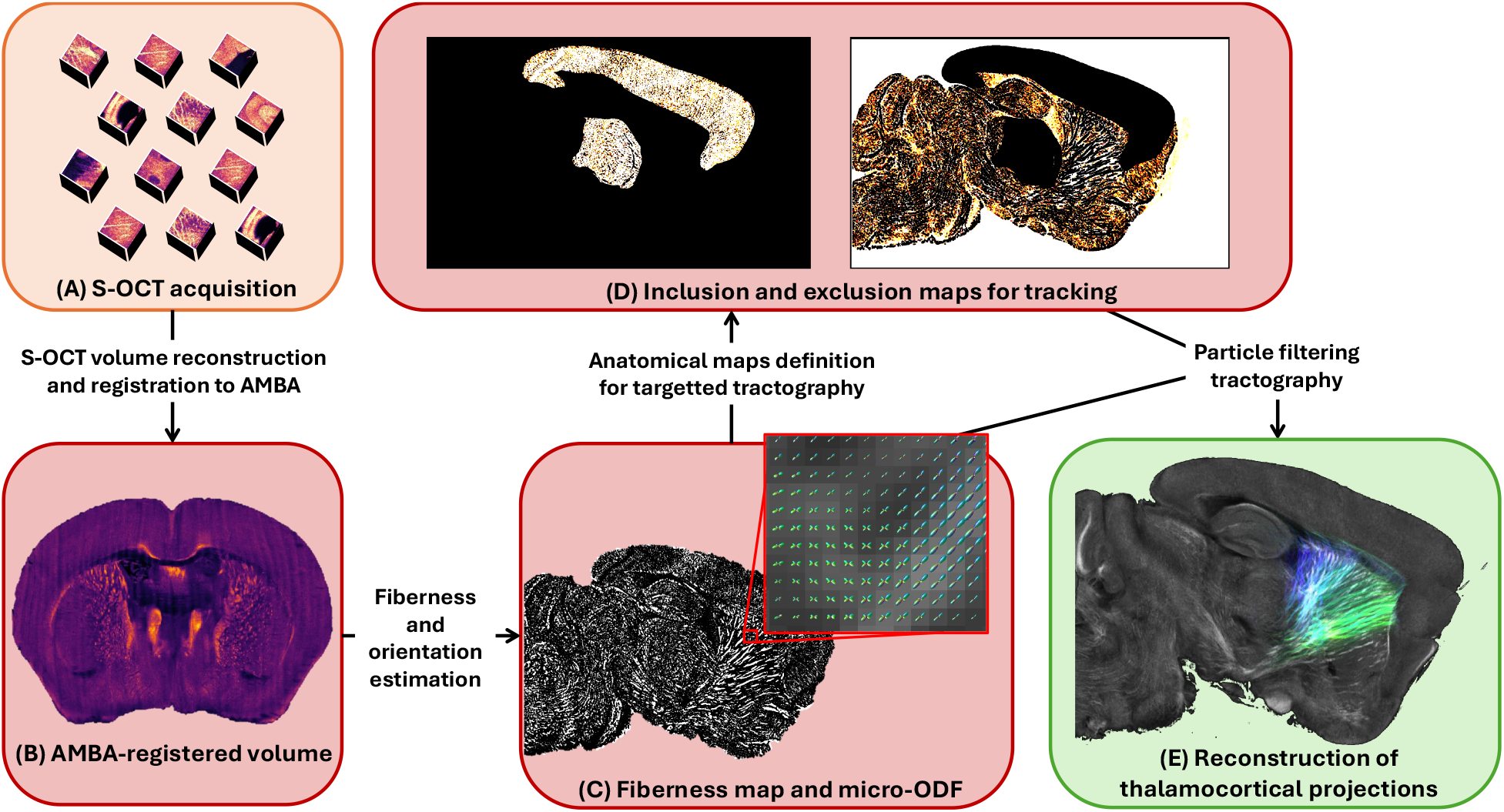
Summary of the steps involved for performing S-OCT tractography. (A) 10000+ volumetric tiles are acquired in serial OCT. (B) These tiles are assembled into a single whole-brain volume and registered to the AMBA. (C) Voxel-wise fiberness and µODF are estimated from the AMBA-registered volume. (D) Inclusion (left) and exclusion (right) maps for tractography are estimated from the fiberness map. (E) A tractography is performed from the inclusion/exclusion maps and µODF.

### 2.1. S-OCT acquisition and volume reconstruction

The S-OCT acquisition used in this work comes from the dataset described in Lefebvre et al. (2024). The Fourier-domain serial OCT (S-OCT) system is equipped with a long working distance objective (M Plan Apo NIR 10X, Mitutoyo, Japan) and has an axial resolution of approximately 3.5 µm and a lateral resolution of around 3 µm. A dissected mouse brain is first mounted onto a motorized stage. Then, volumetric tiles, covering a surface of 750 × 750 µm^2^ (lateral scan) and depth of ≈ 600 µm (axial scan), are acquired sequentially by translating the motorized stage until the entire blockface is imaged. After, a slab of approximately 200 µm thickness is removed from the sample by means of a vibratome. Then, the sample is moved up by 200 µm and the newly revealed blockface is imaged. A typical S-OCT acquisition with 10X objective takes 2-3 days to acquire and results in 10000+ files, amounting to terabytes of raw data. Because many brains from the dataset are incomplete due to errors during acquisition, e.g. out-of-focus samples or tissue tearing, we limit our analysis to a single specimen acquired with its lateral side facing towards the imaging objective, such that the cutting interface is aligned with the sagittal plane. We reconstruct a whole-brain S-OCT volume at a resampled isotropic resolution of 10 µm, using a Nextflow pipeline (Tommaso et al., 2017) built upon the pipeline from Lefebvre et al. (2017, 2018). Reconstruction is done on high-performance computers (HPC) from the Digital Alliance of Canada. The reconstruction pipeline is presented in A. To compensate for tissue deformations, we align the reconstructed S-OCT volume to the Allen Mouse Brain Atlas (AMBA) (Wang et al., 2020), an average mouse brain template built from 1675 mouse brains acquired in serial 2PE fluorescence microscopy and containing annotations for WM and GM structures. We use the 25 µm template for registration. Registration is performed using ANTs (Avants et al., 2011) symmetric diffeomorphic registration. Prior to registration, the S-OCT volume is first converted to nifti and roughly aligned to the AMBA template using ITK-SNAP (Yushkevich et al., 2006). We design a custom objective function for driving deformable registration. This function is the combination of (i) the mutual information between the AMBA and S-OCT volume, (ii) the mutual information between the AMBA annotations and the S-OCT volume and (iii) the cross-correlation between the absolute value of the Laplacians of the AMBA template and the S-OCT volume. This objective function was found to yield satisfying results with our data. Although the optimization is doneat 25-µm resolution,the registered S-OCT volume is kept at 10 µm isotropic resolution to preserve as many detail as possible.

### 2.2. Estimation of fiberness and principal fiber orientation

We use the multiscale method from Sorelli et al. (2023) for estimating the dominant orientation of WM fascicles. This method extends the multiscale vessel enhancement filters from Frangi et al. (1998) and allow for estimating the principal orientation of tubular elements present in an image. Given an image *I*(*x*), we define *I*(*σ, x*) = *G*(*σ, x*) * *I*(*x*) as the scale-space representation of *I*(*x*) at scale *σ*, where *G*(*σ, x*) is a Gaussian filter with standard deviation *σ*, and * is the convolution operator. Let *H* (*σ, x*) be the scale-normalized Hessian matrix of *I* (*σ, x*), with | *λ*_1_ (*σ*) |≤ |*λ*_2_ (*σ*) | ≤ |*λ*_3_ (*σ*)| the sorted absolute eigenvalues of *H* (*σ, x*). From these eigenvalues, we can measure the *blobness R*_*B*_(*σ*), *tubeness R*_*A*_(*σ*) and *structureness S*(*σ*) of the image at voxel position *x*, given by

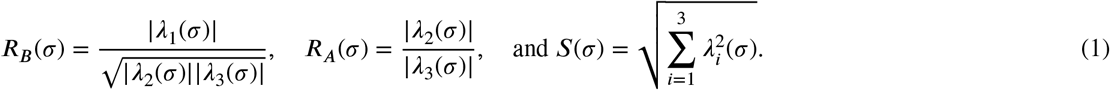

These three metrics are combined into a vesselness score using the equation

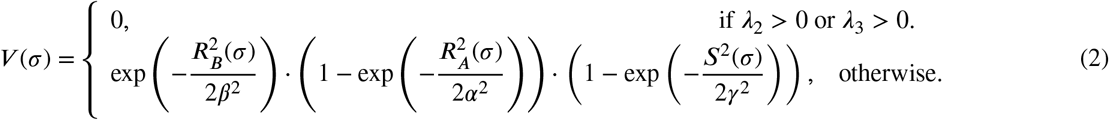

When applied to WM imaging modalities, this vesselness score can be interpreted as the probability of observing a WM fiber, controlled by three parameters α, β and γ. As such, we will refer to this score as the *fiberness* score, going forward. A small value of β will be more agressive in penalizing structures that tend towards a perfect sphere than a high value. Conversely, a small α value will less strongly attenuate the fiberness score as a structure diverge from a perfect tube than a high α value. While these first two parameters depend on the application, the γ parameter is contrast dependent, and is usually set to half the maximum norm of the Hessian matrix *H*(*σ*) (Frangi et al., 1998). In this work, we set α = β = 0.5, as in Frangi et al. (1998), as these values were found to yield satisfying results on our data. The scale-normalized fiberness score *V* (*σ*) can be fairly compared across scales. The fiberness score for some voxel is therefore given by the maximum response of the fiberness filter, i.e.

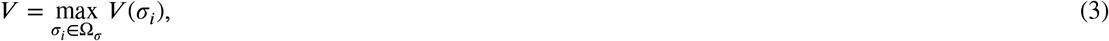

where Ω_*σ*_ is the set of all evaluated scales. Then, the orientation of a structure at voxel *x* is given by the eigenvector *v*_1_ corresponding to eigenvalue *λ*_1_, i.e. the direction of weakest intensity variation.

### 2.3. Estimation of a microscale ODF

As in Sorelli et al. (2023), we use an analytical method (Alimi et al., 2020) to estimate a microscale orientation distribution function (µODF) from the estimated local orientations. The orientation (*θ*_*x*_, *ϕ*_*x*_) for voxel *x* can be represented as a Dirac delta function of the form

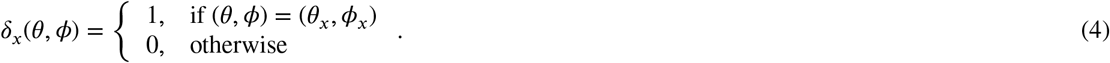

In dMRI tractography, spherical functions are often expressed as a spherical harmonics (SH) series (Tournier et al., 2007; Descoteaux et al., 2007). With *Y*_*l,m*_(*θ, ϕ*) the real and symmetric SH function of even order *l* ≥ 0 and degree *m* (™*l* ≤ *m* ≤ *l*), the Dirac delta function describing the orientation (*θ*_*x*_, *ϕ*_*x*_) at some voxel *x* is given by the SH series

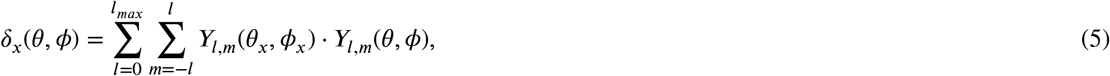

with *l*_*max*_ is the maximum SH order, set to 8 in this work, and *Y*_*l,m*_(*θ*_*x*_, *ϕ*_*x*_) is the SH function evaluated along orientation (*θ*_*x*_, *ϕ*_*x*_).

To mitigate Gibbs ringing and negative-amplitude lobes arising from truncating the SH delta function to *l*_*max*_, we also incorporate apodized Dirac delta functions. This approach is also implemented in Zhang et al. (2025), where structure tensor orientations are expressed as apodized Dirac delta functions before summing them over the super-voxel. The apodized Dirac delta function is obtained from an optimization routine penalizing for negative or small positive lobes resulting from Gibbs ringing. For more details on the algorithm, the reader is referred to (Raffelt et al., 2012; Dhollander et al., 2014). The result of the optimization routine is a spherical convolution kernel *R* that can be applied to the original Dirac delta function:

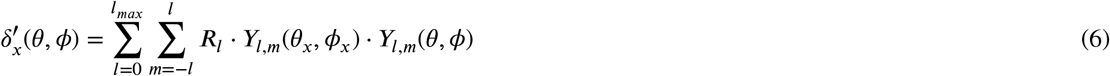

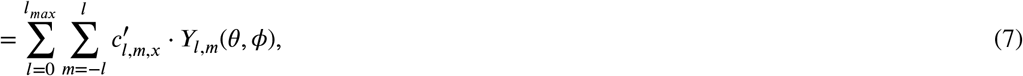

with 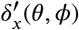 the apodized Dirac delta function for orientation (*θ*_*x*_, *ϕ*_*x*_) and 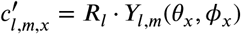 the multiplicative scalar ^*x*^SH coefficient at voxel *x* for order *l* and degree *m*.

In Alimi et al. (2020), an ODF is estimated inside a super-voxel covering *K* voxels by summing up the contribution of all orientations expressed as Dirac delta functions in SH coefficients. This super-voxel approach is often implemented by downsampling the voxel grid and pooling together neighbouring voxels to create a µODF (Alimi et al., 2020; Sorelli et al., 2023, 2025). However, this approach would constrain S-OCT tractography to the coarser resolution of the resampled voxel grid. Here, we instead use a uniform sliding window formulation (as in (Zhang et al., 2025)) which is evaluated for each native voxel position *x*, given by

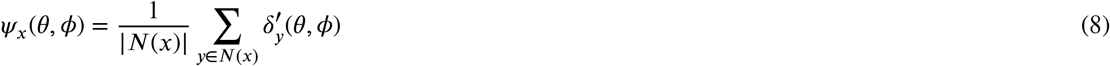

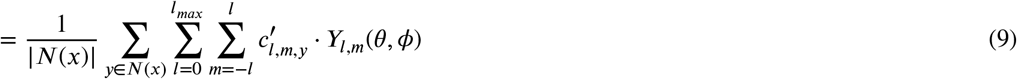

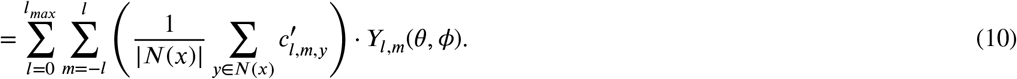

Here, *N*(*x*) is the set of voxels located inside a *ω* × *ω* × *ω* voxels^3^ centered on, and *N*(*x*) denotes the cardinality of the set. The term between parentheses is therefore a scalar coefficient *c*_*l,m,x*_ multiplying the SH function *Y*_*l,m*_(*θ, ϕ*). With 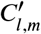 a coefficients array such that 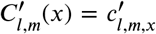, the coefficients array *C*_*l,m*_(*x*) for the resulting µODF can be obtained via convolution, i.e.

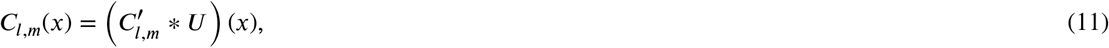

where * describes the convolution operoator and *U* (*x*) is a uniform filter of width *ω* given by

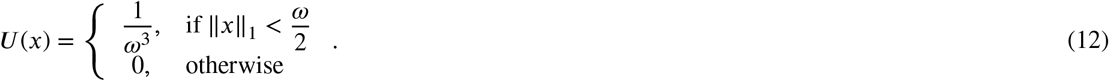

### 2.4. Particle filtering tractography and anatomical maps definition

We define probabilistic maps for tractography using the continuous maps criterion (CMC) (Girard et al., 2014). Instead of applying thresholds to tissue partial volume effects (PVE) maps, as in anatomically-constrained tractography (Smith et al., 2012), CMC employs a probabilistic approach to determine the status of a streamline. CMC relies on two tracking maps: (i) Map^in^ describes the probability of a streamline being considered valid (included) based on where it is terminated; (ii) Map^ex^ describes the probability of a streamline being considered invalid (excluded) based where it is terminated. For human brain dMRI tractography, Map^in^ is usually defined as the GM PVE map to include streamlines terminating in GM, and Map^ex^, as the cerebrospinal fluid (CSF) PVE map to discard streamlines terminating inside ventricles voxels. The CMC maps are evaluated at each tracking step to determine whether a streamline can continue. For a given voxel position *x*, the probability of a streamline continuing its propagation is given by

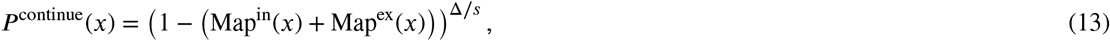

where Δ is the voxel size and *s* is the step size. Hence, as the probability of being either included or excluded increases, the probability of continuing decreases. When a streamline fails the propagation test, the probability of it being included in the final tractogram is given by

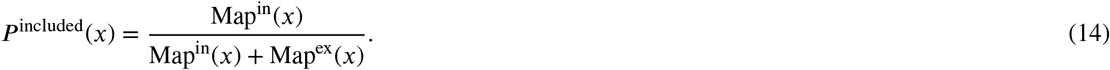

CMC is used jointly with particle filtering tractography (Girard et al., 2014) (PFT), a tracking module for handling backtracking. Backtracking is a mecanism that has been first described in (Smith et al., 2012) for retrying tractography a few steps before a streamline has failed an inclusion test, in the hope that drawing a new sample at each step will result in a valid trajectory. PFT estimates many streamline trajectories from a prior position along the streamline, located at a backward distance of *δ*_*b*_ mm from the termination event. Each candidate trajectory is propagated along a forward distance of *δ*_*f*_ mm. Then, a trajectory is drawn from the set of valid trajectories, and streamline propagation resumes. The number of trajectories sampled during backtracking is controlled by the particles count, which should be set high enough to be representative of the underlying distribution, but as small as possible to keep computation requirements low (Girard et al., 2014). For human brain dMRI tractography, the particles count is set to 15. If no valid trajectory is found, the whole streamline is excluded from the output. Backtracking is only allowed for a fixed number of trials, such that a streamline reaching an invalid region doesn’t get stuck in an infinite loop.

GM makes for most of the mouse brain. As such, for mouse brain S-OCT tractography, long-reaching WM connections are unlikely to be reconstructed as often as short connections, as the probability of leaving WM and reaching a valid termination region increases with the number of steps taken. As such, we modify CMC to consider the targeted tractography problem, where we attempt to reconstruct WM paths between pairs of specific anatomical regions, instead of any GM-GM connection. Then, with ℛ the set of voxels where streamlines are allowed to terminate, we update the definitions for Map^in^ and Map^ex^ such that

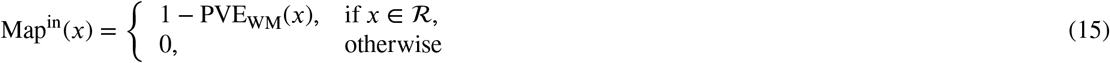

and

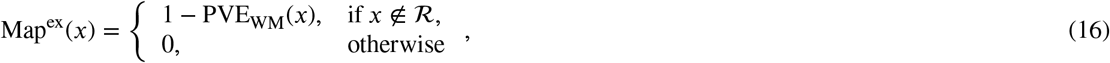

where PVE_WM_ is the WM probability at voxel *x*. For diffusion MRI tractography, PVE maps are often estimated using segmentation algorithms applied to T1-weighted images, which are unavailable for S-OCT. Here, we use the fiberness measure from Equation 3 to estimate the probability of a voxel belonging to WM. To increase WM probability inside low contrast regions, we first clip the fiberness between 0 and the 95^th^ percentile before rescaling to the range [0, 1]. We then use the rescaled fiberness map as the WM partial volume effect map PVE_WM_. To remove holes in the tracking maps that may arise from S-OCT acquisition and reconstruction artifacts, we apply a grayscale erosion to Map^th^ using a 3-voxel-wide box as the structuring element.

## 3. Experiments

This work comprises two complementary evaluations. First, a simulated microscopy-like phantom with known ground truth is used to compare the proposed µODF estimation method and tractography algorithm to commonly employed alternatives. Second, our tailored approach is applied as a proof of concept to a single whole mouse-brain S-OCT dataset acquired in the sagittal plane. The real-data experiment is intended to assess feasibility and anatomical plausibility rather than population-level reproducibility or comprehensive anatomical validity.

### 3.1. Quantifying microscale tractography with simulated data

#### 3.1.1. Simulated dataset creation

We generate a microscopy-like dataset from the ground-truth (GT) streamlines of the simulated Fibercup phantom (Fillard et al., 2011) from Neher et al. (2017). The objective of the simulation dataset is to evaluate how well a tractography algorithm is able to reconstruct the trajectories of individual WM fascicles. This differs from the traditional dMRI tractography validation approaches that reconstruct macroscale WM bundles (Côté et al., 2013; Maier-Hein et al., 2017; Renauld et al., 2023). We first downsample the 7 GT bundles from the Fibercup dataset to 25% their initial number of streamlines, and then reconstruct track-density images (TDI) (Calamante et al., 2012) of each bundle at 250 µm isotropic resolution using tckmap from MRtrix (Tournier et al., 2019). These values (25% and 250 µm) were chosen experimentally such that the number of GT streamlines inside each voxel is close to 1 and that there is empty space between the simulated WM fascicles. To simulate the contrasts observed in OCT images, the value of each voxel is increased by 0.4 and multiplicative noise is simulated by drawing samples from a gamma distribution (as speckle patterns in OCT have been shown to follow a gamma distribution (Kirillin et al., 2014)), resulting in a signal-to-noise ratio (SNR) of 100. This SNR was chosen based on the SNR observed from real S-OCT reconstructions at 10 µm isotropic resolution. The resulting simulated dataset is shown in Figure 3 (B.). Because the bundles from the FiberCup phantom are aligned on a single plane, the resulting contrast is representative of in-plane WM fascicles measured in S-OCT. We also estimate track-orientation density images (TODI) (Dhollander et al., 2014) from the downsampled GT streamlines using MRtrix, from which we extract the GT fiber orientations at each voxel using `scil_fodf_metrics` from scilpy (Renauld et al., 2026). The GT orientations inside two regions of interest are shown in Figure 3 (C.-D.).

**Figure 3.**
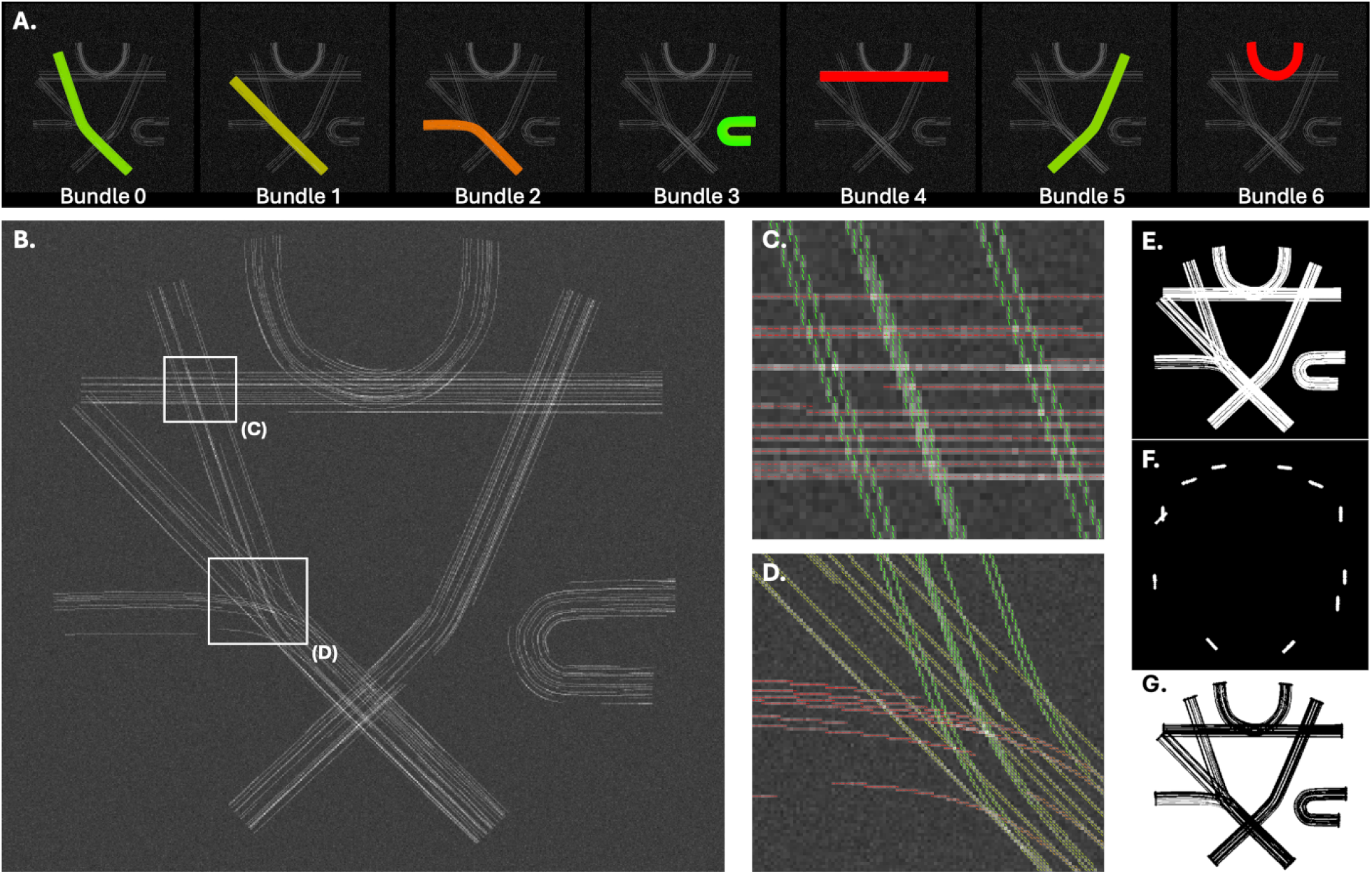
Simulated dataset overview. A track-density image (B.) is generated at 250µm isotropic resolution from the downsampled ground-truth bundles (A.). To simulate the contrasts observed in OCT images, the value of each voxel is increased by 0.4 and speckle noise is simulated by drawing samples from a gamma distribution. Subfigures (C.) and (D.) show ground-truth fiber orientations at each voxel for the regions framed in (B.). The tracking mask for local tracking, corresponding to the dilated WM mask (TDI > 0), is shown in (E.) and the inclusion and exclusion maps are shown in (F.) and (G.).

#### 3.1.2. Quantitative evaluation of µODF estimation

We then compare the µODF estimation method from Sorelli et al. (2023) to structure tensor analysis. The structure tensor method is described in B and depends on a single scale parameter ρ. We estimate principal orientations from multiscale Frangi filters for scales *σ* ∈ {0.5; 1.0; 1.5} voxels, and for scales ρ = 0.5, ρ = 1.0 and ρ = 1.5 voxels for structure tensor analysis. We also study the effect of the 3-dimensional spatial averaging window width for combining individual voxel orientations into a µODF for widths of 7, 11 and 15 voxels. To assess the quality of the µODF estimation, we compare it to the ground-truth orientations and report the voxel-wise maximum angular error. For each ground-truth orientation, the angular error is computed as the minimum angle among all peaks extracted from the estimated µODF. If no peaks are extracted, a maximal error of 90^°^ is assigned. Then, the voxel-wise maximum angular error, corresponding to the peak most poorly aligned out of all the ground-truth peaks inside a voxel, is estimated.

#### 3.1.3. Quantitative evaluation of tractography reconstruction

We quantify our tractography reconstructions using the Tractometer (Côté et al., 2013; Maier-Hein et al., 2017; Renauld et al., 2023), implemented in scilpy (Renauld et al., 2026). The GT bundles masks are obtained by thresholding the noise-free TDI map for each bundle. We also create endpoint masks for the head and tail of each bundle, which we then dilate 10 times using a 3-voxel wide structure element. These pairs of endpoints masks are used to find valid streamlines, i.e. streamlines connecting two valid endpoints. To discard implausible streamlines that would go beyond the expected extent of a bundle, we generate additional inclusion masks by applying a gaussian blur (*σ* = 1) to the noiseless bundle TDI map, which we then threshold. This results in hole-free bundle inclusion masks, and a streamline belonging to a bundle must fit entirely within this mask to be considered valid. We compare PFT to deterministic and probabilistic local tractography. We use the same tracking parameters for all methods: step size of 0.5 voxel, maximum angle of 20 degrees between two consecutive steps, and relative threshold on ODF amplitudes of 0.1. The WM mask, obtained by the union of all GT bundle masks, is used for seeding, using 5 seeds per voxel (npv = 5). The same seeding positions are used throughout all experiments. For local deterministic and probabilistic tractography, we use a once-dilated WM mask as our tracking mask (Figure 3, E.). Inclusion and exclusion maps for PFT are generated from the WM and endpoints masks. The inclusion map is defined as the union of all bundles heads and tails, dilated 5 times (F.). The exclusion map is defined as the inverse of the union of the one-time dilated WM mask and inclusion map (G.). The forward and backward tracking distances for PFT are kept to default values of *δ*_*f*_ = 1 mm and *δ*_*b*_ = 2 mm. We also compare tractography on µODF estimated using the Frangi filters method to those estimated from the structure tensor method. A description of the reappraised Tractometer scoring system used in this work can be found in Renauld et al. (2023) and useful metrics for this work are summarized in C.

### 3.2. Experiments with real S-OCT data

#### 3.2.1. S-OCT tractography

We demonstrate S-OCT tractography using the AMBA-aligned S-OCT reconstruction at 10-µm isotropic resolution described in subsection 2.1. As a reminder, this specimen was acquired with the cutting interface is aligned with the sagittal plane. The registered S-OCT volume is shown in Figure 4. The optimal scale for detecting some structure should be set to half its expected radius (Sorelli et al., 2023). Visual analysis of the S-OCT dataset revealed that WM fascicles have a radius between 10 µm to 40 µm (1 to 4 voxels). As such, the scales used for multiscale Frangi filters were set to *σ* ∈ {0.5; 1.0; 1.5; 2.0} voxels. A small region of interest of 100 × 100 × 100 voxels (1 mm^3^; identified by white box in Figure 4) covering the anterior portion of the CC and a portion of the internal capsule is chosen for selecting the optimal spatial averaging window width among values 7, 11 and 15. The spatial averaging window width for estimating µODF is determined qualitatively by visually comparing the reconstructed µODF to the underlying S-OCT contrast. Orientations are then estimated for the whole brain at 10 µm isotropic resolution, and combined into µODF using the filter width found from previous step.

**Figure 4.**
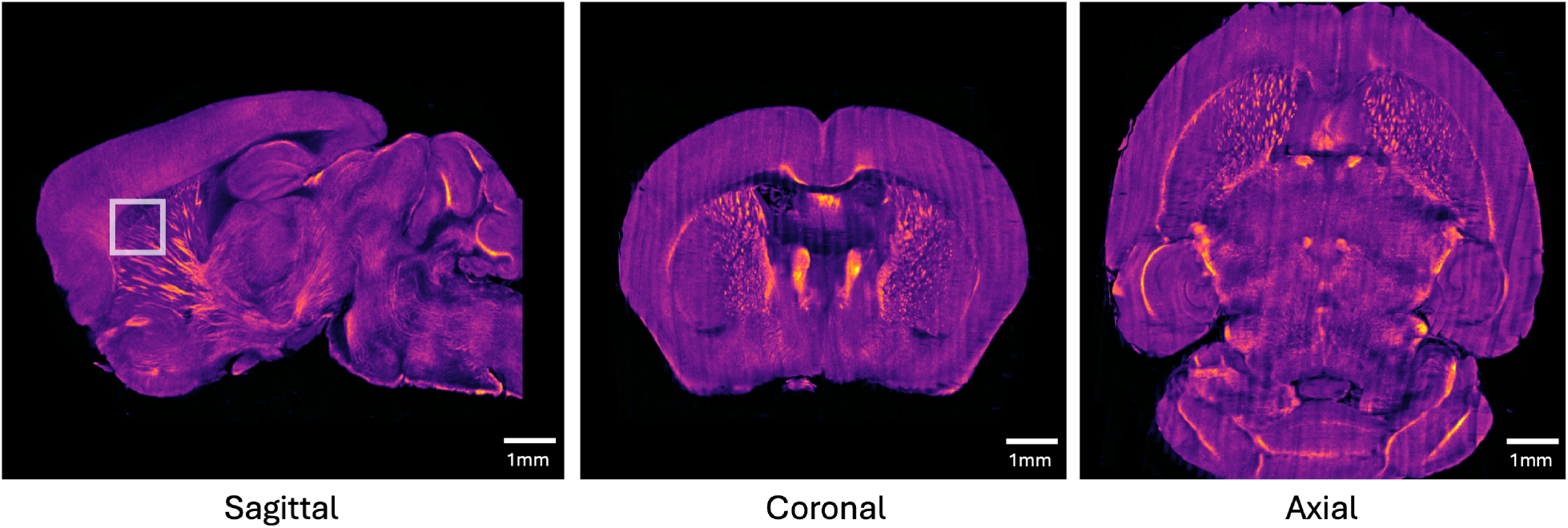
S-OCT reconstruction at 10µm isotropic voxel size after registration to the Allen Mouse Brain Atlas (AMBA) along the 3 standard views. The sagittal view corresponds to the plane of acquisition. The region of interest for parameter search is identified by the white frame.

For tractography, we consider the problem of reconstructing WM fascicles projecting from the thalamus to the isocortex. A schematic representation of the problem is shown in Figure 5 (top row). WM fascicles connecting the isocortex to the thalamus (green dotted lines) must cross the internal capsule, a WM structure hidden inside the caudoputamen (CP). While a non-targeted, whole-brain tractography approach would consider any streamline terminating inside the CP (bottom row, white) as a valid streamline, here we only allow streamlines to terminate either along the isocortex (bottom row, blue) or inside the thalamus (bottom row, green). The seeding mask is defined as voxels inside the CP with a non-zero fiberness. We use brain segmentations from the AMBA to identify the thalamus, isocortex and CP. We split the µODF volume and CMC maps into a left and right hemisphere. This allows for reducing the memory requirements for tractography, while also preventing the reconstruction of commissural connections. To determine the optimal tracking parameters for our S-OCT data, we first perform a grid search on step sizes (Δ ∈ {0.002, 0.005, 0.010} µm) and maximum curvature angle (*θ* ∈ {20, 40, 60} degrees). We also evaluate the effect of varying the particles count for PFT between 15 (default for human dMRI tractography), 50 and 100 particles. For each parameter combination, we track from the same 100,000 seeding positions, randomly sampled from the seeding mask. The resulting streamlines are filtered such that all valid streamlines have a single endpoint in the cortex and a single endpoint in the thalamus. This parameter search is done for the left hemisphere only. We define the optimal tracking parameters as the ones maximizing the number of streamlines successfully connecting the isocortex and the thalamus. PFT forward and backward tracking distances are scaled with respect to the voxel size such that *δ*_*f*_ = 40 µm and *δ*_*b*_ = 80 µm. We set the minimum streamline length to 400 µm.

**Figure 5.**
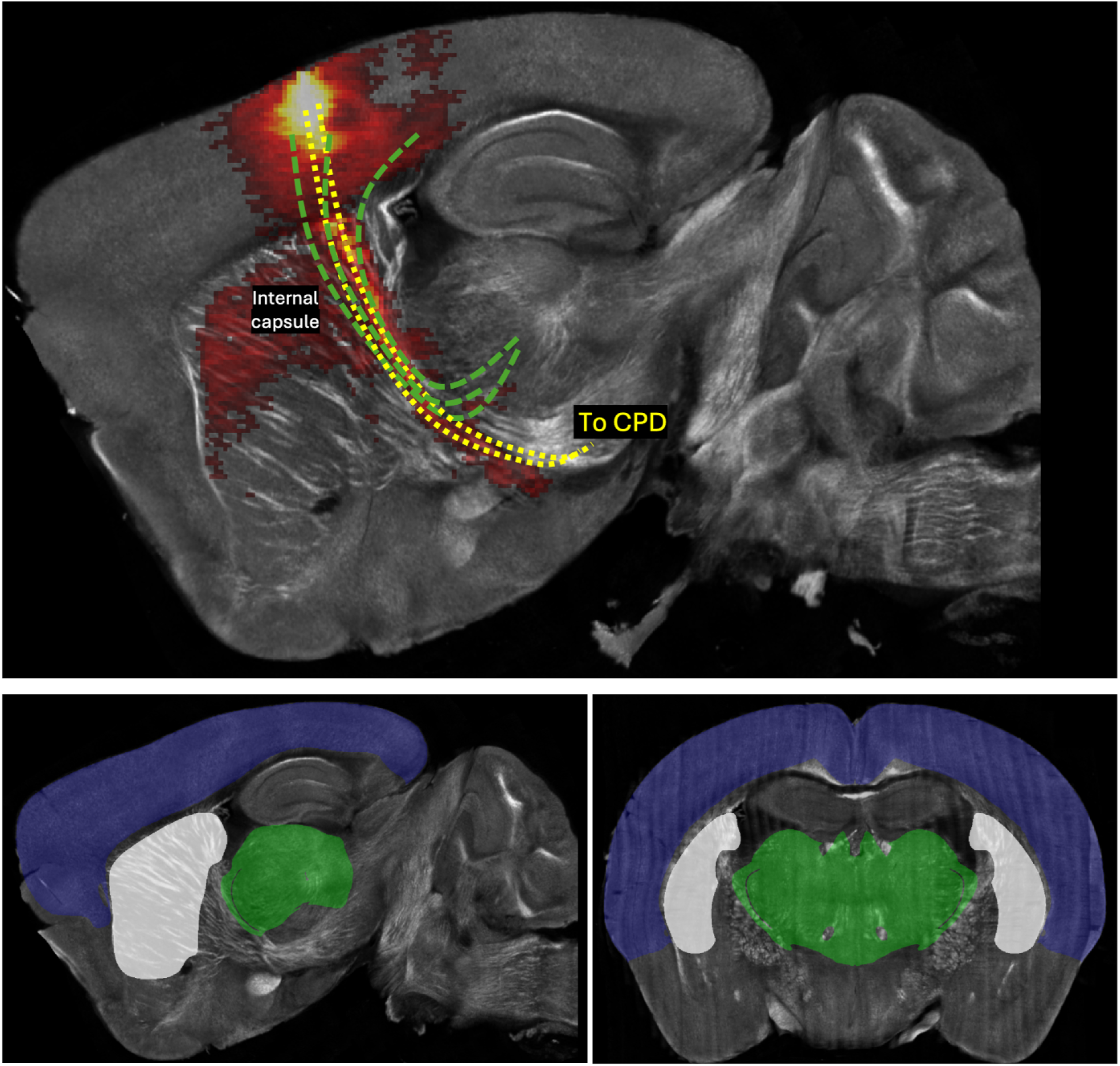
Reconstructing thalamocortical projections from S-OCT. *Top:* Schematic description of thalamocortical (green dotted lines) and corticospinal (yellow dotted line) projections visible from an S-OCT acquisition. The heatmap shown in overlay shows a viral tracer experiment from the Allen Mouse Brain Connectivity Atlas highlighting these projections. *Bottom:* Brain segmentations for CMC maps definitions, in sagittal (left) and coronal (right) views. The isocortex is shown in blue, the thalamus, in green, and caudoputamen, in white.

Using the optimal tracking parameters found at the previous step, we then run PFT for both hemispheres from 2,000,000 seeding positions. Due to the very high memory and time requirements, we run tractography on clusters from the Digital Alliance of Canada as 20 batches of 100,000 seeds per hemisphere. The resulting tractograms are concatenated into a single tractogram and filtered such that remaining streamlines connect the thalamus to the isocortex.

#### 3.2.2. Comparison with viral tracer experiments

We compare the resulting tractograms to tracer experiments from the Allen Mouse Brain Connectivity Atlas (AMBCA). We first download the injection and projection density maps at 50 µm resolution for all experiments injected in the isocortex. We then binarize the maps by apply a 0.05 threshold. Then, we filter each experiment to keep only those making an intersection with the caudoputamen and thalamus. Because more AMBCA experiments are available for the right hemisphere, we limit our analysis to right thalamocortical projections. Of the 1338 AMBCA experiments with injections in the isocortex, 709 fulfilled these conditions. Then, for each valid experiments, we filter the S-OCT tractography reconstruction for streamlines ending inside the injection mask and fitting entirely within the binarized projection map. To account for intersubject variability and registration errors, we allow a 1-voxel tolerance, i.e. streamlines coordinates can diverge from the expected binary masks if they are within 1 voxel of the mask.

### 3.3. Data and code availability statement

The simulated FiberCup dataset and derivatives archived on Zenodo (Poirier, 2026b). The real S-OCT dataset and derivatives are also available on Zenodo as a separate archive (Poirier, 2026a). Because of the 50GB size limit, whole-brain µODF are not included in the archive, but are available upon request. Processing was done using opensource softwares. In particular, linumpy^1^ (v0.0.1) was used for reconstructing the S-OCT volumes. ANTs (Tustison et al., 2021) was used for registration (Avants et al., 2008). We integrated the code for multiscale Frangi filters^2^ into the scilpy toolbox (Renauld et al., 2026), which we used for estimating µODF, for tractography and for scoring the tractograms. The meso2macro (m2m) toolkit (Abou-Hamdan et al., 2023) was used for downloading the average brain and viral tracer experiments from the Allen atlas.

### 3.4. Ethics statement

No data was acquired exclusively for the purpose of this work. Data from Lefebvre et al. (2024) was approved by the Animal Research Ethics Committee at the Montreal Heart Institute (protocol 2023-23-05).

## 4. Results

### 4.1 Fibercup µODF estimation

The distribution of voxel-wise maximum angular errors between the principal fiber orientations extracted from the estimated µODF and the ground-truth fiber orientations is shown in Figure 6a (top row) for the Frangi method (*σ* ∈ {0.5; 1.0; 1.5} voxels). Results are reported for spatial averaging kernel widths *ω* ∈ {7; 11; 15} voxels. We report the distribution of angular errors inside single-fiber voxels (NuFO = 1, black), and crossing voxels (NuFO = 2, blue).

**Figure 6.**
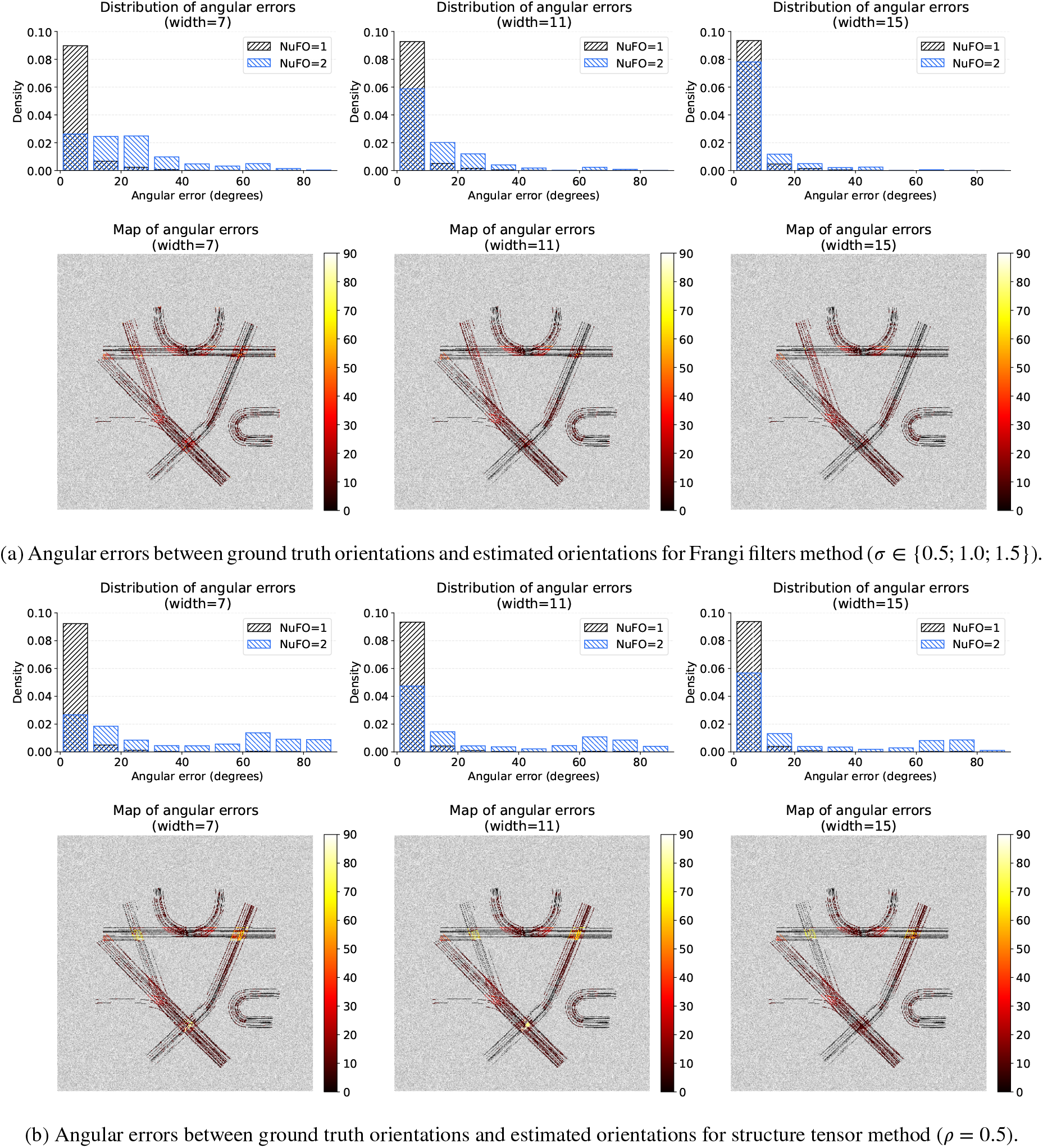
Distributions of angular errors between ground truth orientations and estimated orientations for Frangi and structure tensor methods for varying spatial averaging window width (*ω*).

Results for the structure tensor method are shown in Figure 6b (top row). The reported values for structure tensor analysis are shown for optimal scale parameter ρ = 0.5, resulting in the lowest angular errors. For structure tensor analysis, we note that a poorly-tuned scale parameter ρ results in an increase in angular errors (result not shown). As such, we expect that the multiscale Frangi filter method is more reliable for real data, where WM fascicles vary in diameter. For single-fiber voxels, the distribution of angular errors remains similar for all *ω* values for both methods. However, inside crossing-fibers voxels, we report a higher density of voxels with high angular errors for the structure tensor method than for the Frangi filter method. In bottom row of both subfigures, the angular errors are shown as a heatmap, overlayed on the simulated contrast. The highest errors are observed in crossing regions for all methods. Errors close to 90^°^ are likely due to a single principal orientation being extracted inside a 2-fibers voxel.

The mean maximum angular errors for both methods are reported in Table 1 for *ω* ∈ {7; 11; 15}. For an equivalent spatial averaging window width, the Frangi filter method systematically results in lower errors for NuFO = 1 and NuFO = 2. The method resulting in the lowest voxel-wise maximum angular error is the Frangi filter method with *ω* = 15 with an average maximum angular error of 3.91^°^ inside NuFO = 1 voxels and 10.40^°^ inside NuFO = 2.

**Table 1.** Mean maximum angular errors for Frangi filters and structure tensor method for varying spatial averaging window width. Values are reported for NuFO = 1 and NuFO = 2.

| Method | Width | Mean error<br>(NuFO = 1) | Mean error<br>(NuFO = 2) |
| --- | --- | --- | --- |
| Structure tensor | 7 | 5.25° | 36.73° |
|  | 11 | 4.81° | 28.11° |
|  | 15 | 4.59° | 23.18° |
| Frangi filters | 7 | 5.50° | 23.63° |
|  | 11 | 4.25° | 14.43° |
|  | 15 | 3.91° | 10.40° |

### 4.2. Fibercup tractography

Tractometer scores for deterministic, probabilistic and particle filtering tractography (PFT) are shown in Table 2 for the Frangi filter method, for *ω* ∈ {7; 11; 15}. The reported metrics are averaged over all 7 bundles. Deterministic tractography is the method resulting in the highest number of reconstructed streamlines (count column), with nearly twice as many streamlines compared to the other methods. Only 2 valid bundles (VB) could be recovered using probabilistic tractography, with less than 50 valid streamlines (VS) reconstructed. Although deterministic tractography is the method with the highest streamlines count, it could only recover 5 out of the 7 ground-truth bundles, with a VS ratio of 5.19% – 5.72%. PFT is the only method recovering all 7 VB. Despite having the lowest streamlines count, PFT achieves the highest VS count and VS ratio. Increasing the spatial averaging window width *ω* decreases the mean Dice coefficient (F1) and mean overlap (OL). The highest-scoring tractography reconstruction with respect to mean Dice coefficient (F1) is obtained with PFT and *ω* = 7 (F1 score of 38.81%). This low F1 score is explained by the high bundle overreach (mean OR): 74.45% of the reconstructed VB extend beyond the boundaries of the GT bundle masks.

**Table 2.**
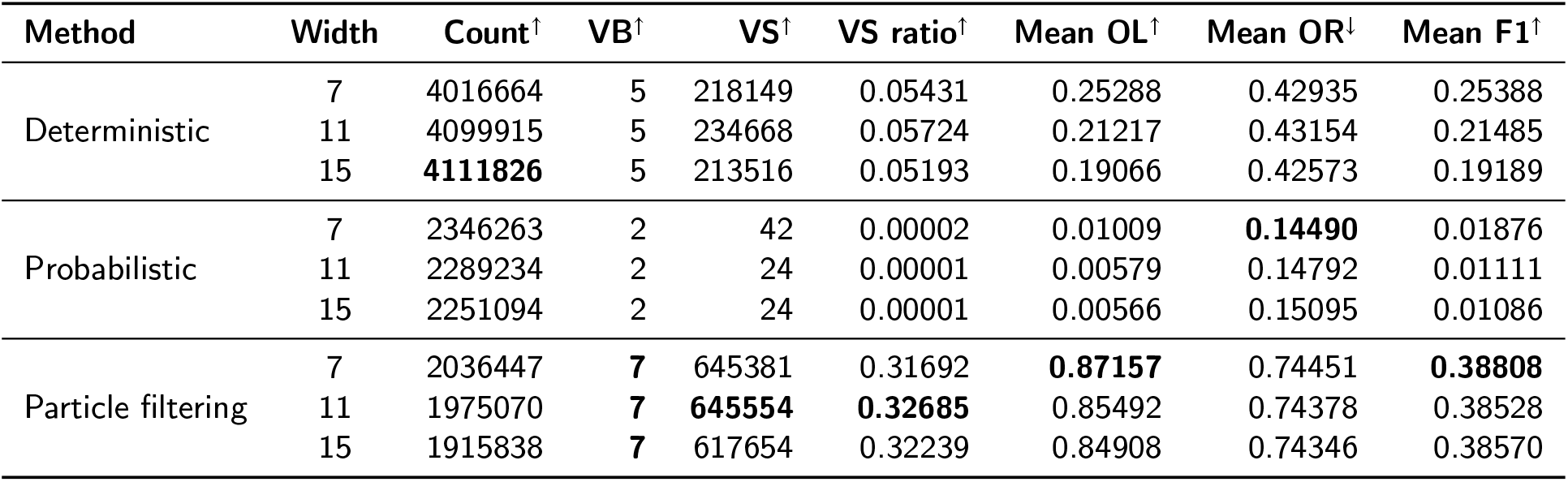
Tractometer metrics for the simulated S-OCT contrast for the Fibercup dataset for deterministic, probabilistic and PFT from Frangi filter micro-ODF. The optimal value for each metric is shown in bold.

The bundle-wise OL, OR and VS count are reported in Figure 7 for spatial averaging window widths *ω* ∈ {7; 11; 15}. We report high overlaps for all bundles, except bundle 0 (39.75%). The reconstructed fiber bundles are shown in Figure 8, overlayed on the simulated Fibercup contrast. Problematic fiber configurations intersected by each bundle, e.g. crossing, splitting or kissing configurations, are identified by white circles. For hard-to-track bundles such as bundle 0 and bundle 2, making a 3-way splitting with bundle 1, using a smaller *ω* allows for better specificity to the underlying WM fascicles, resulting in an improved overlap compared to higher *ω*. Bundle 1 sits roughly midway between bundles 0 and 2 and scores better then both other bundles and is less sensitive to the choice of spatial filtering window width. Based on OL and VS count, bundle 0 is the hardest to reconstruct: in addition to being the bundle with the highest number of crossings, bundle 0 also makes the smallest crossing angle with bundle 1. As we see in Figure 8, almost no VS cover the inferior-left portion of bundle 0, where it overlaps with bundle 1 and 2. The bundle-wise overreach (OR) plot shows that all bundles overreach to a similar extent. We also note that changing *ω* has little effect on the reported OR. Bundle 3 and bundle 6 maximize VS and OL. From Figure 8, we see that these are the two bundles intersecting the lowest number of problematic regions. The remaining bundles 1, 2, and 5 all cross as many problematic regions (2), and have a similar VS count.

**Figure 7.**
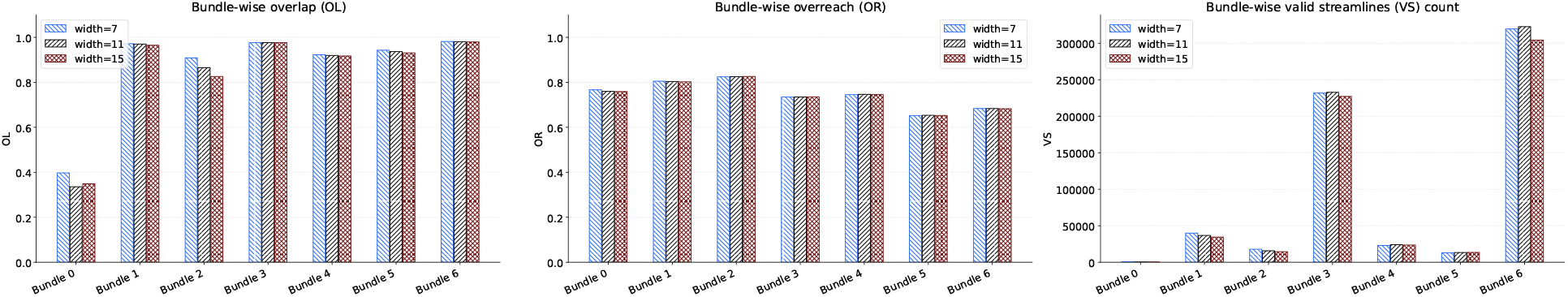
Effect of width of mean filter on overlap, overreach and VS count, per bundle. Bundle 0 is very poorly reconstructed compared to other bundles. For all bundles except bundle 0 and bundle 2, increasing the width of the filter has very little impact on thereconstruction metrics.

**Figure 8.**
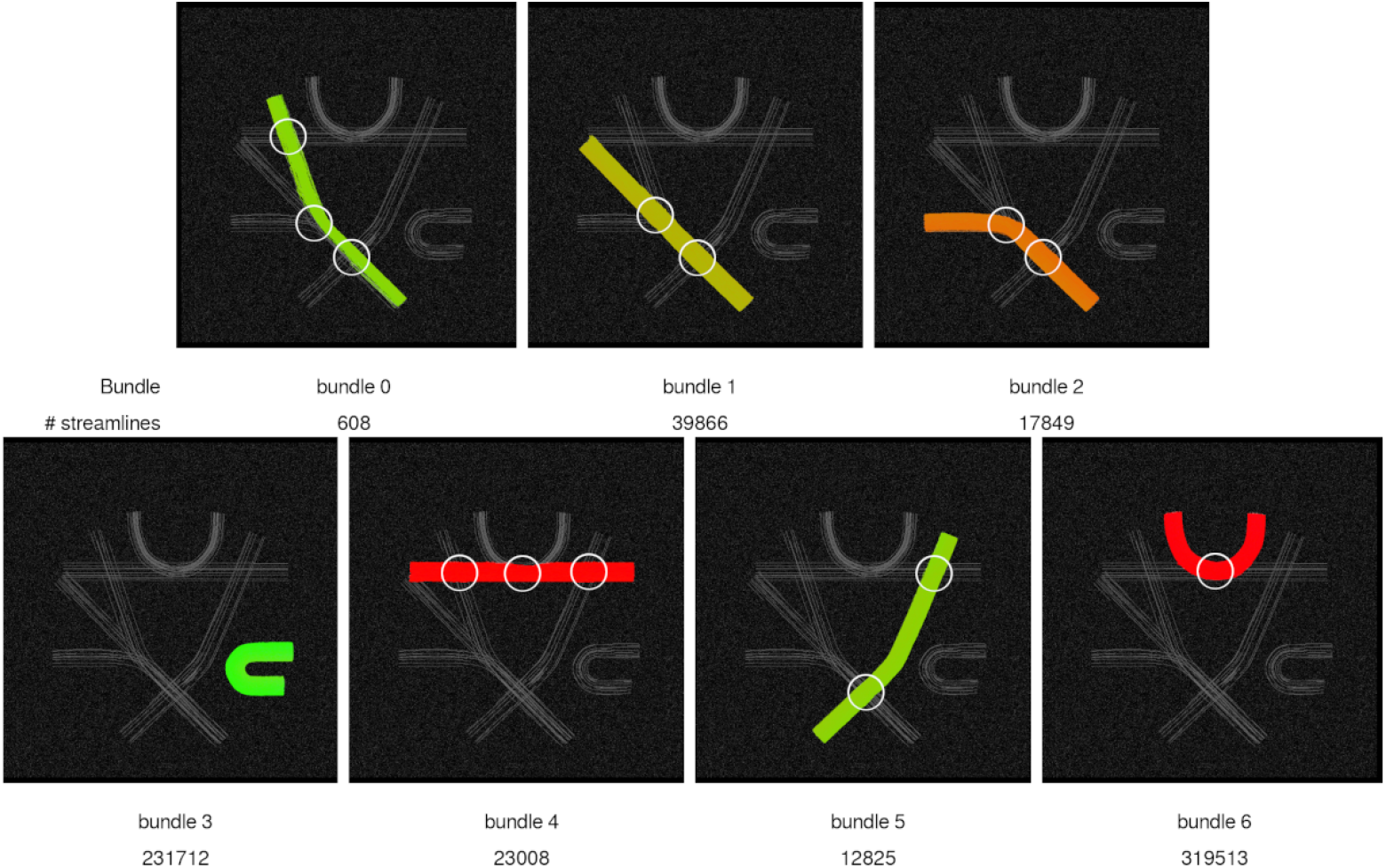
Valid bundles recovered using particle filtering tractography with Frangi filter µODF for spatial averaging window width *ω* = 7. The number of valid streamlines (VS) recovered for each bundle is reported below. The circles highlight regions with problematic fiber configurations intersected by each bundle, such as crossing, splitting and kissing configurations.

### 4.3. S-OCT tractography

The estimation of µODF was done on high-performance computers (HPC) from the Digital Alliance of Canada. The 10-µm resolution S-OCT volume registered to the AMBA is saved as a 2.5 GB nifti file. Then, estimating the principal fiber orientation per voxel required 316.6 GB of random access memory (RAM) and 4.6 hours on a single CPU. The resulting fiber orientations volume is 3.1 GB in size. µODF estimation required 550.6 GB of RAM for 4.7 hours and resulted in a 77 GB SH volume (maximum order 8). We note that Sorelli et al. (2025) have described a multinode version and a GPU-accelerated version of their method, both of which we did not try. This choice is motivated by a desire to keep requirements minimal such that the shared computational resources from the Digital Alliance of Canada are more easily usable. Splitting in two hemispheres resulted in two 39 GB volumes. Each tracking experiment required 505.6 GB of RAM and took between 1 and 4 hours, depending on the tracking parameters and seeding positions.

Figure 9 shows the tracking maps estimated using our approach combining annotations from the AMBA with the fiberness measure from the multiscale Frangi filters. The fiberness map, in (B.), is maximal for in-plane WM fascicles. The royal blue circle highlights WM fascicles showing up faintly in the original S-OCT image, but which have a strong WM probability according to the fiberness measure. The multiscale nature of the fiberness measure allows for identifying WM fascicles of different scales. The cyan circle shows WM fascicles located inside the CP and surrounded by GM, where we see that WM fibers are assigned a high WM probability while surrounding GM is assigned a low WM probability. The fiberness measure is only sensitive to in-plane WM fibers, assigning a low probability to out-of-plane WM structures, e.g. the fimbria (yellow) or the anterior forceps of the CC (green). The inclusion and exclusion maps are shown in (C.) and (D.), respectively. The probability of inclusion is high for streamlines terminating along the isocortex and thalamus, and null for all other brain regions.

**Figure 9.**
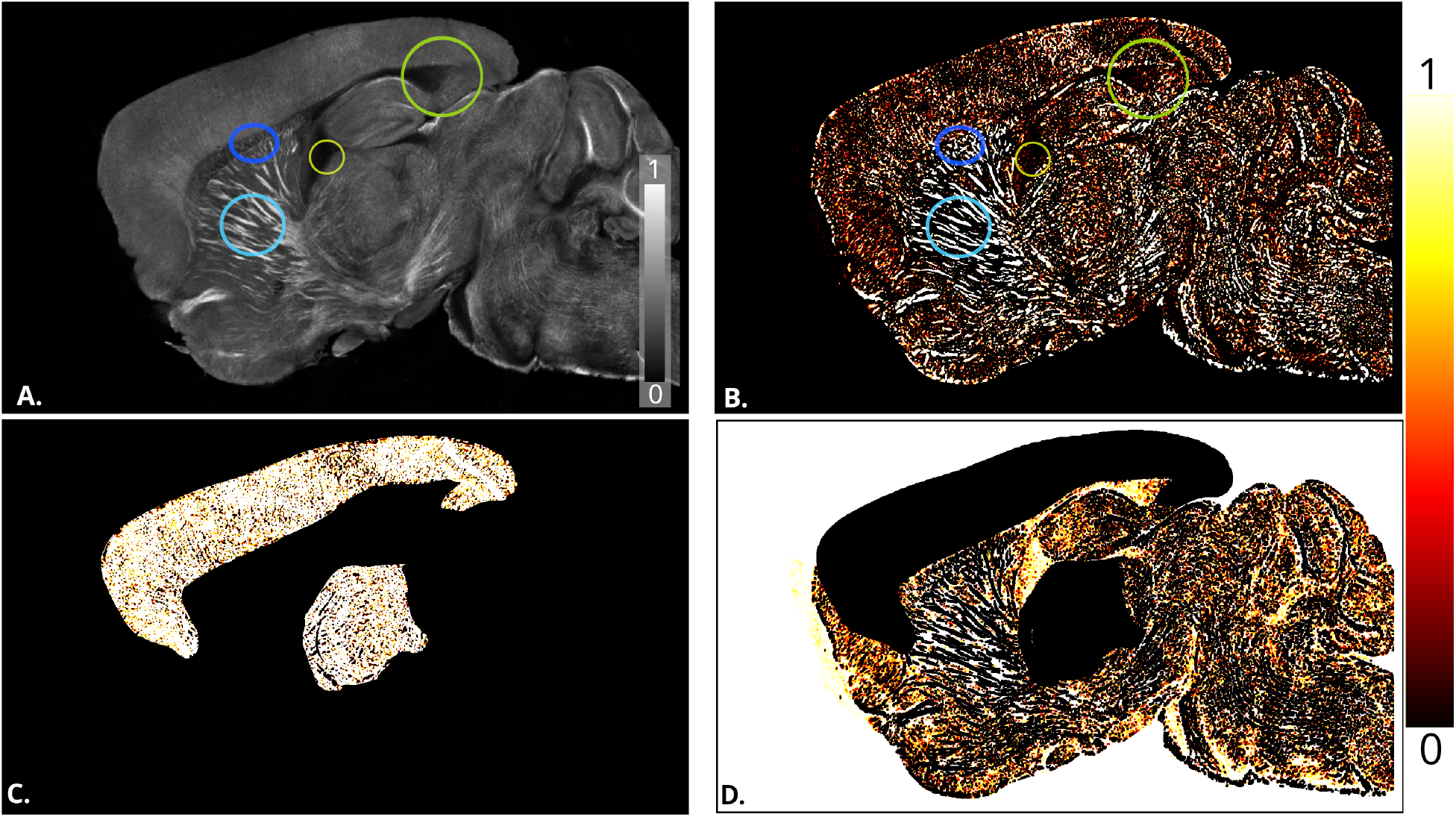
Anatomical maps estimated from S-OCT using Frangi filter. Annotated circles are described in the main text. **A.**S-OCT volume. **B**. Fiberness measured with Frangi filters, thresholded to 95^th^ percentile and rescaled to the range [0, 1]. **C**. Inclusion map. Only streamlines terminating inside the isocortex or thamamus have a high probability of being included in the final tractogram. **D**. Exclusion map. Streamlines reaching regions other than the isocortex or thalamus will be terminated.

The µODF estimated from our mouse specimen are shown in Figure 10 for spatial filtering window widths *ω* ∈ {7; 11; 15}, inside a small crop of 100 × 100 × 100 voxels (red patch, subfigure A.). The cropped region is enlarged in (B.). Subfigures (C1.) to (C3.) show the µODF estimated inside the red frame. We see that using a small filtering window of width 7 better recovers crossing µODF (highlighted by red arrow, enlarged in E.) than higher values. Increasing *ω* smoothes the µODF field and decreases the specificity of µODF to the underlying WM structures, as shown in subfigures (D1.) to (D3.) for the yellow frame in (B.). A low *ω* constrains the estimated µODF to the underlying fascicles, allowing interstitial GM space between WM fascicles to remain free of µODF (yellow arrow, enlarged in F.). Based on these conclusions, the spatial filtering window width selected for estimating µODF for the whole brain was set to *ω* = 7 voxels, to remain specific to the underlying WM fascicles while recovering crossing fiber configurations in complex fiber regions.

**Figure 10.**
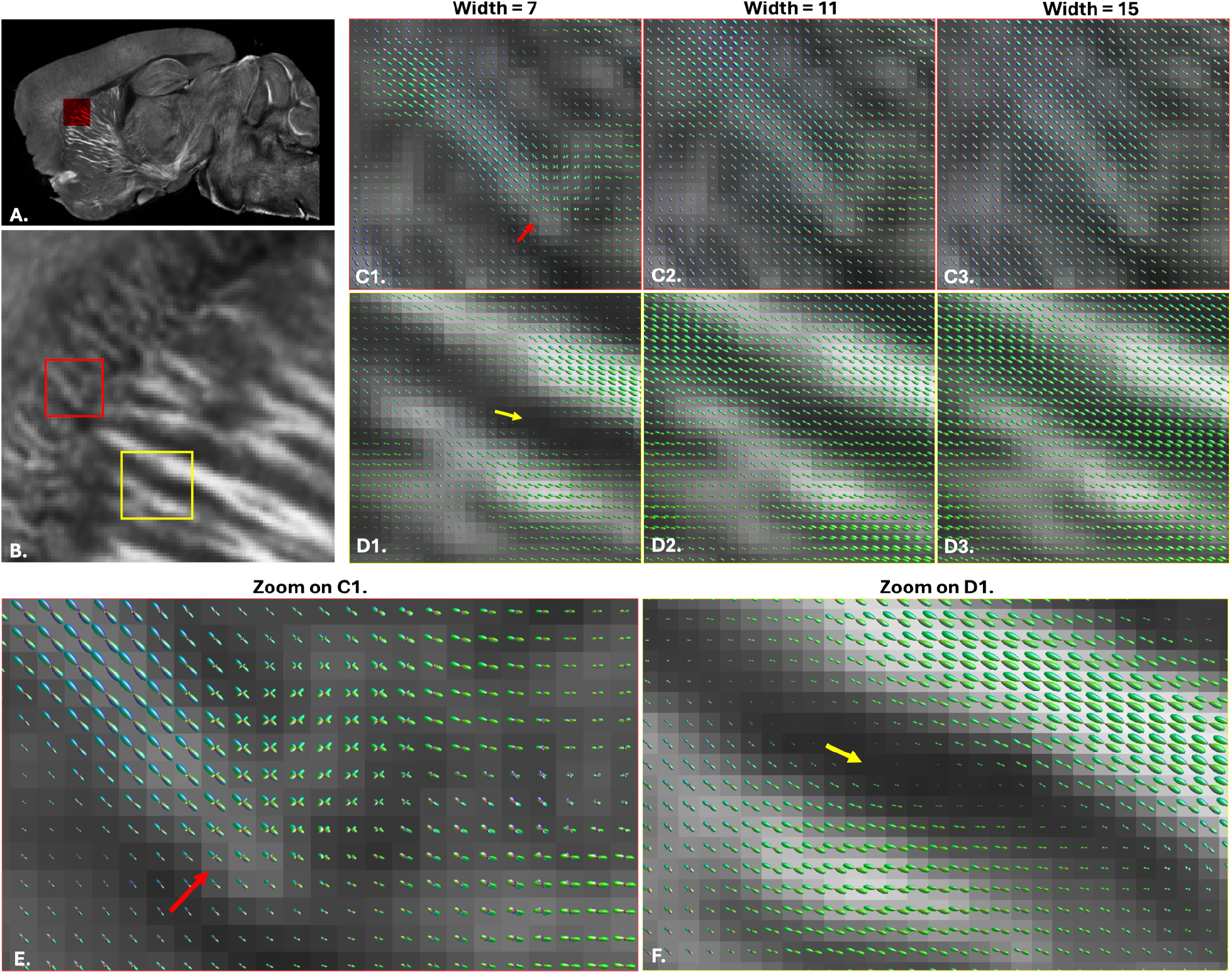
µODF estimated inside a small crop from our S-OCT specimen at 10 µm isotropic resolution for varying spatial averaging window width (*ω*). *A*. Whole-brain S-OCT reconstruction, with the red box representing the crop considered for parameter search. *B*. Zoom of the cropped volume, with two regions of interest identified by the red and yellow frames. *C*. µODF estimated inside the red region for increasing *ω*. *D*. µODF estimated inside the yellow region for increasing *ω E*. Zoom on C1. *F*. Zoom on D1.

In Figure 11, we report the number of streamlines obtained using PFT for different combinations of step size Δ ∈ {0.002, 0.005, 0.010} µm, *θ* ∈ {20, 40, 60} degrees and particles count ∈ {15, 50, 100}. For each parameter combination, PFT was run from the same 100,000 seeds and the resulting tractogram was filtered such that the remaining streamlines have one endpoint in the isocortex and one endpoint in the thalamus. Parameter search was done for the left hemisphere only. Small step sizes result in few valid streamlines compared to bigger step sizes. For an equivalent *θ*, a smaller step size results in a smaller radius of curvature, meaning streamlines are more likely to deviate from the forward propagation direction. The figure shows that parameters Δ = 0.010 µm and *θ* = 40^°^ maximize streamline count, and that increasing the particles count increases the number of streamlines. As such, we set the tracking parameters to Δ = 0.010 µm, *θ* = 40^°^ and particles count = 100 for the rest of this work.

**Figure 11.**
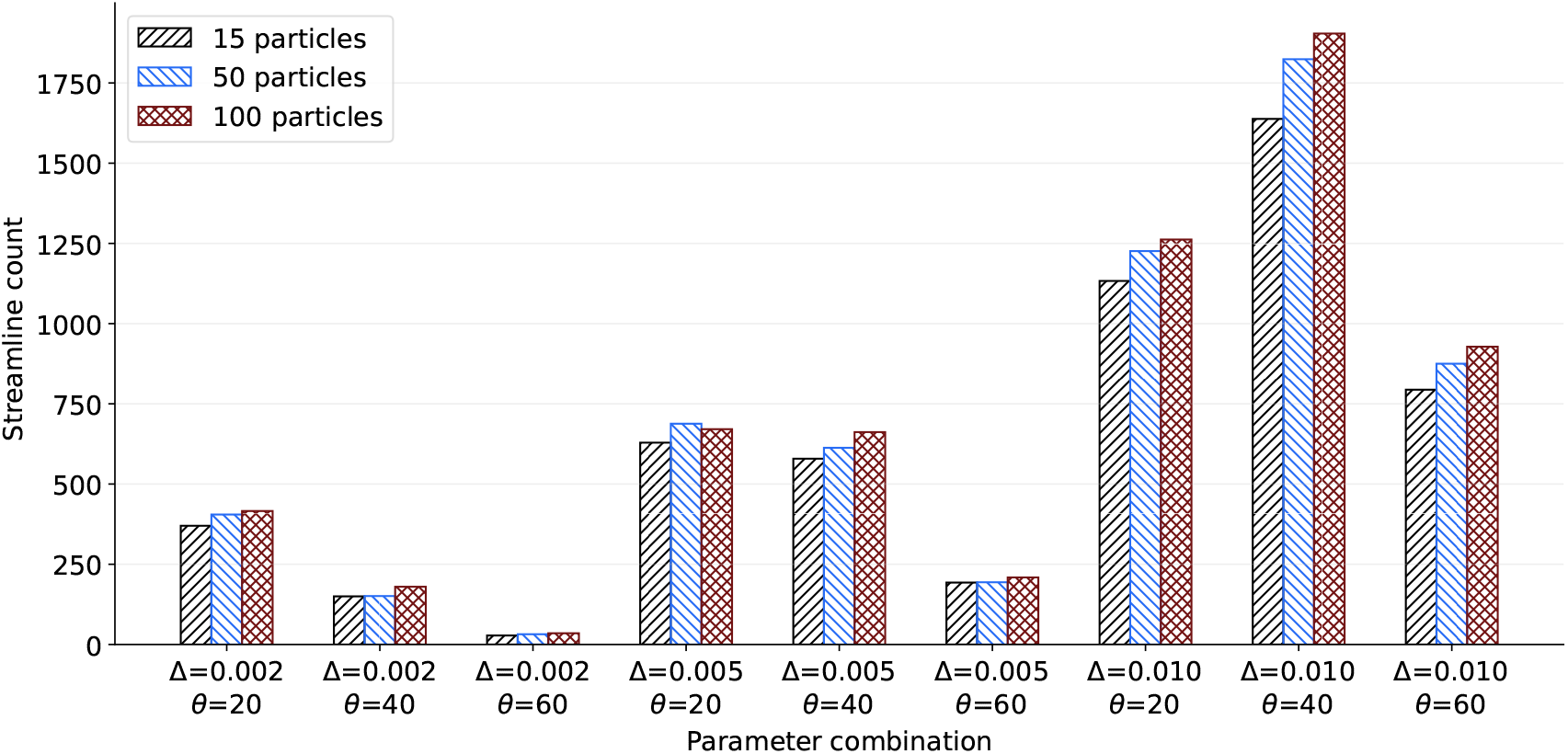
Streamline count for different combinations of step sizes (Δ) and maximum curvature angle (*θ*) and for varying number of particles.

Figure 12 shows the thalamocortical projections reconstructed from S-OCT tractography for the left and right hemispheres, using the tracking parameters found above, after filtering the tractography outputs such that only streamlines connecting the thalamus and isocortex are kept. Tractography is done from 2,000,000 seeding positions per hemisphere, randomly sampled from the seeding mask. After filtering, we obtain 37,498 streamlines for the left hemisphere and 32,286 streamlines for the right hemisphere. Streamlines are organized into clusters overlapping with the underlying WM fascicles visible on the S-OCT volume. Streamlines also successfully follow WM strands appearing weakly on the S-OCT image as they cross the CC. S-OCT tractography successfully reconstructs WM fascicles of different diameters, but bigger fascicles are more densily reconstructed than finer ones, as is expected when using a WM seeding strategy (bigger bundles are sampled more often than finer ones). We also note that S-OCT tractography results in false negatives, with some finer WM fascicles visible on the S-OCT volume not being reconstructed with tractography.

**Figure 12.**
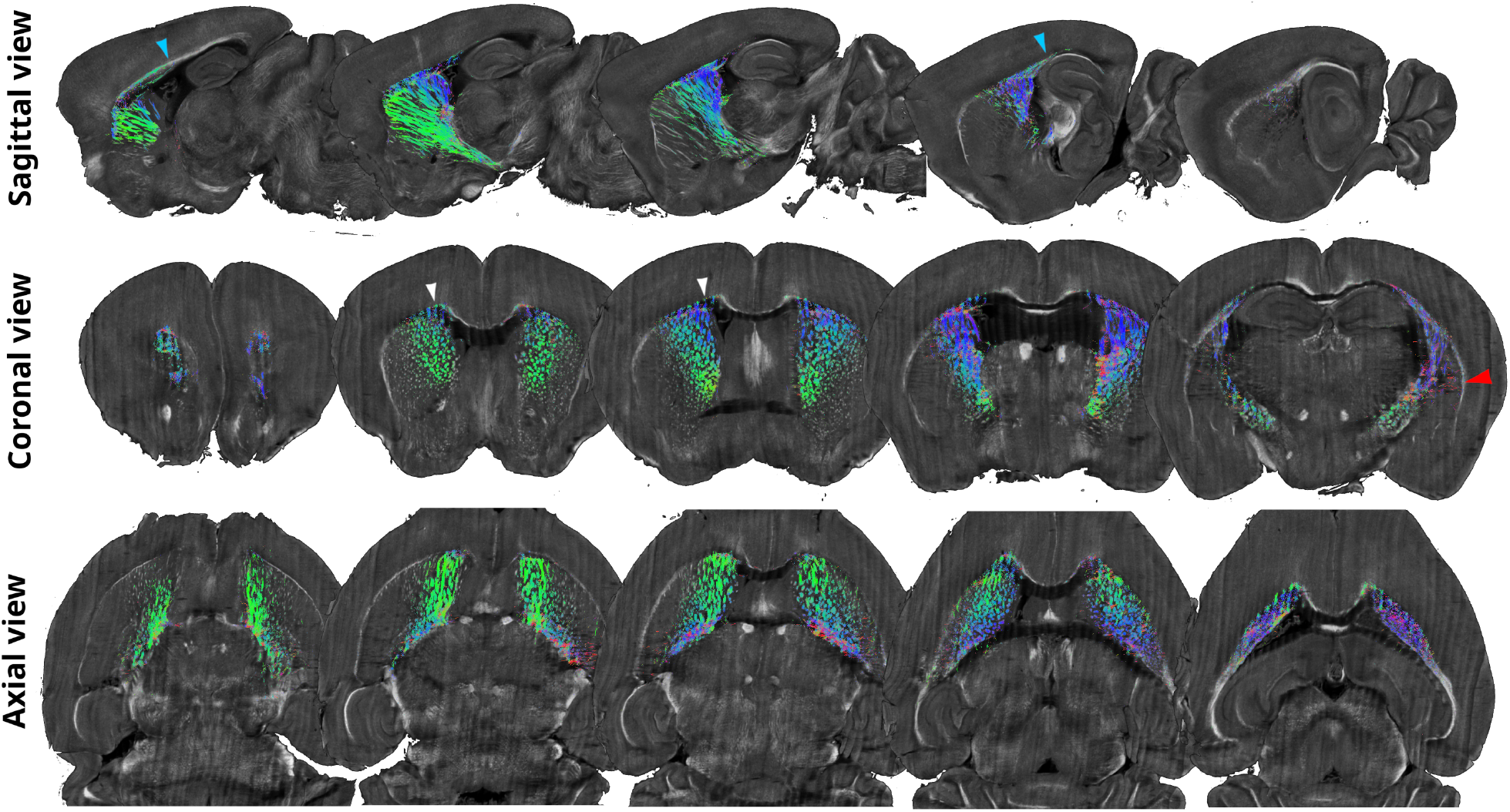
Thalamocortical projections reconstructed from S-OCT tractography for both hemispheres in sagittal, coronal and axial views. The reconstructed streamlines are organized in clusters, aligned with the underlying WM fascicles visible with S-OCT.

The S-OCT volume shows striping patterns along the left-right slicing direction (see coronal and axial views). These stripes align with the surface of the sample blockface, where the signal is higher, and more streamlines are reconstructed along these stripes than in-between (white arrows, coronal view). This is also demonstrated in Figure 13a, where we report the number of streamline points (y-axis) along each sagittal slice (x-axis). The barplot is overlayed onto an example slice in coronal view, and shows that the distribution of streamline points exhibits local maxima aligned with the stripe artifacts. The red arrow in Figure 12 shows that S-OCT tractography reconstructs a few out-of-plane WM connections, although S-OCT is only sensitive to in-plane WM fascicles. In Figure 13b we report the distribution of deviations (in 10-degrees increments) between the sagittal plane and the orientation of streamline segments. We see that streamlines are mostly aligned with the sagittal plane, with a very low proportion of streamline segments having an out-of-plane (90^°^ degrees) orientation. We also note that some streamlines propagate along other WM structures (such as the cingulum) or along the interface between WM and GM (e.g. the superior portion of the CC) instead of directly reaching for GM (cyan arrows, sagittal view).

**Figure 13.**
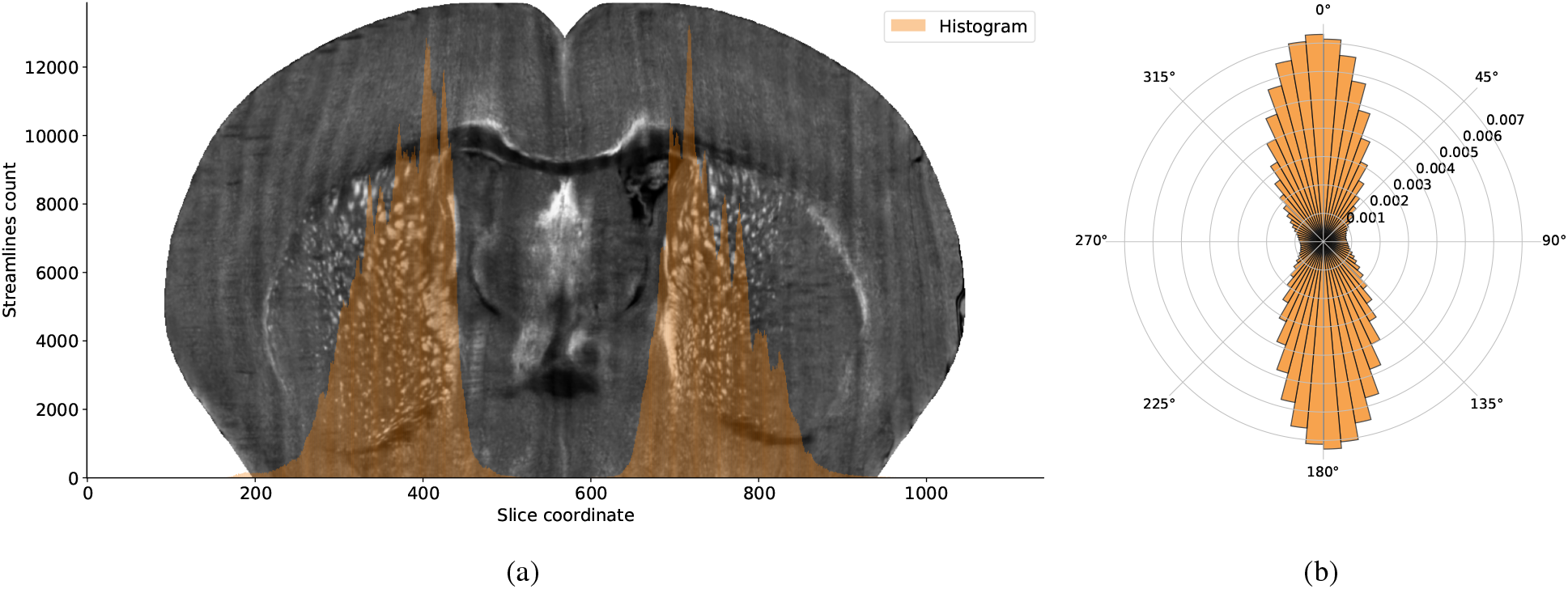
Distributions of streamlines positions and orientations with respect to the acquisition plane. (a) Distribution of streamline coordinates along the left-right axis. The histogram (orange) is overlayed on a coronal slice. (b) Normalized density distribution of deviations (in degrees) between the orientations of streamline segments and the sagittal plane.

Figure 14 shows example viral tracing experiments from the AMBCA which are reproduced from S-OCT tractography. AMBCA projection density maps are shown as a heatmap overlayed on the S-OCT volume. The thalamus is shown in green. Streamlines were filtered such that they end within 1 voxel of the binarized injection site and that all streamlines point are at a distance smaller than 1 voxel from the binarized projection density map. Out of the 32,286 streamlines reconstructed for the right hemisphere, 11,635 streamlines were reproducible from viral tracer injections (36% valid streamlines ratio). This value is close to the one reported on simulated data (33%). We however note that it is possible that the remaining “invalid” streamlines were simply not covered by any experiment from the AMBCA. It could also be that the streamlines deviate slightly from the expected tracer trajectory, which may be due to intersubject variability and not anatomical implausibility. The viral tracing experiments most often reproduced by S-OCT tractography are those injected in the primary somatosensory (SSp) area, primary somatomotor area (MOp), and orbital area (ORBl), all of which are reachable by propagating along the plane of acquisition.

**Figure 14.**
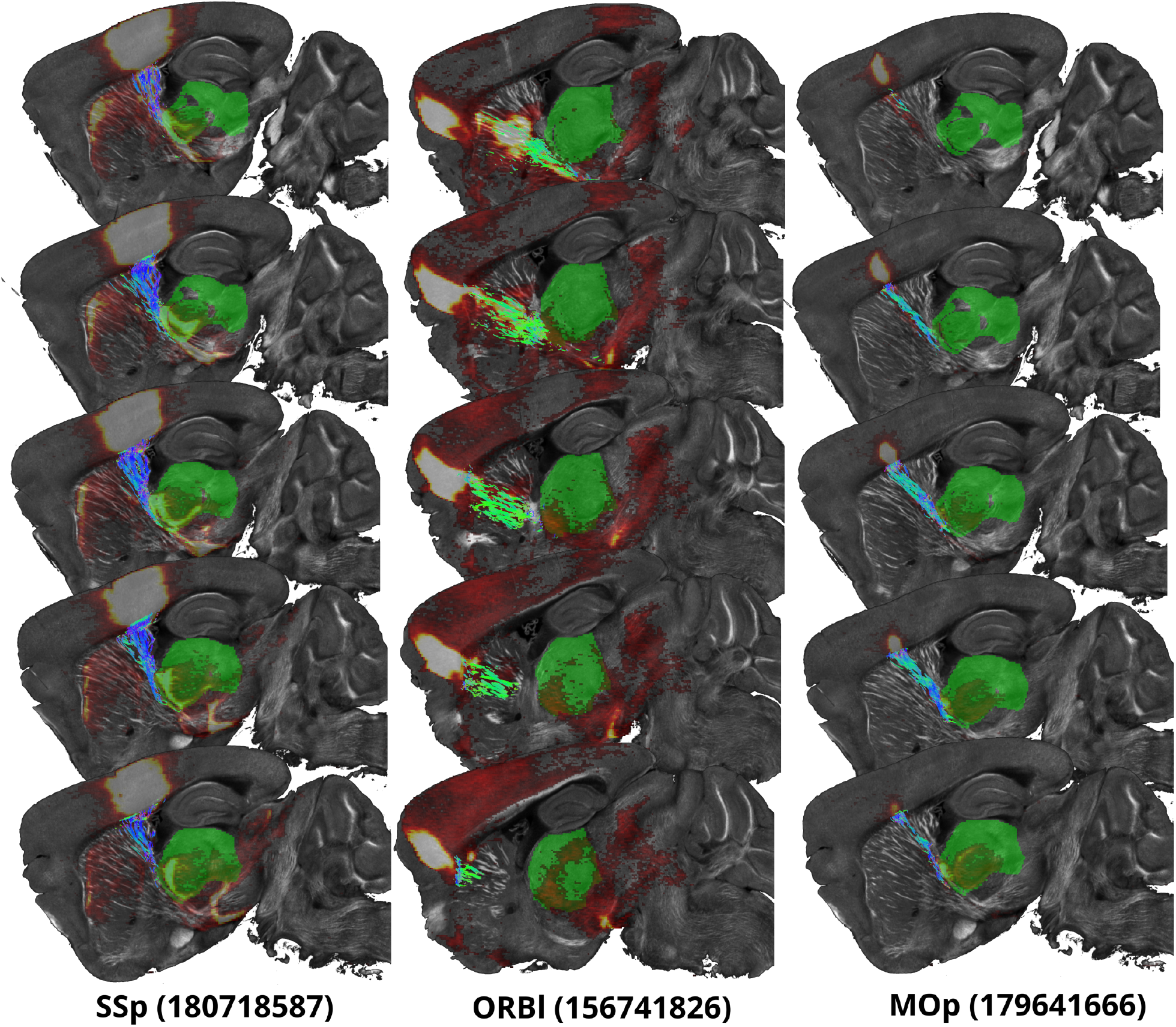
Example thalamocortical projections obtained from S-OCT tractography reproducing viral tracing experiments from the AMBCA for primary somatosensory area (SSp), primary motor area (MOp) and orbital area, lateral part (ORBl). The streamlines (directional-encoding colormap) and viral tracer projection maps (hot colormap) are overlayed on the S-OCT volume. The thalamus is shown in green. The identifiers for each of the experiments are written in parentheses. Experiments can be visualized on https://connectivity.brain-map.org/.

## 5. Discussion

In this work, we described a tailored approach for tractography from serial optical coherence tomography acquisitions. Using simulated data, we showed that going beyond the structure tensor, commonly used for microscopy tractography, in favor of a multiscale, higher-order Frangi filter method (Sorelli et al., 2023) improves the quality of the estimated µODF. We also described a continuous method for reconstructing the µODF at native resolution, by replacing the discrete super-voxel approach by a sliding window formulation (Zhang et al., 2025). We further built upon the work of Sorelli et al. (2023) by using apodized Dirac delta functions for representing fiber orientations in spherical harmonics coordinates. We also showed that, in the presence of noise and ambiguous fiber configurations, the combination of anatomical constraints and probabilistic particle filtering tractography (PFT) outperforms both deterministic and probabilistic local tractography approaches for S-OCT tractography. We also showed that the fiberness maps obtained from multiscale Frangi filters can be used for estimating anatomical tissue maps from a whole mouse brain S-OCT reconstruction, enabling targeted anatomically-constrained tractography. We demonstrated our approach on an S-OCT volume at 10 µm isotropic resolution for reconstructing thalamocortical WM projections visible on S-OCT acquisitions. Using a 10-µm S-OCT reconstruction, we showed that PFT successfully reconstructs thalamocortical WM projections visible on S-OCT acquisitions. We further evaluated the anatomical accuracy of the reconstructed streamlines by comparing them to viral tracing experiments from the Allen Mouse Brain Connectivity Atlas and showed that S-OCT tractography reconstructs anatomically-plausible WM connections.

### S-OCT tractography: How?

#### Frangi filter over structure tensor

We used simulated data to assess the quality of orientation estimations and tractography from S-OCT volumes. To assess the quality of the reconstructed µODF, we measured the maximum angular error with the ground-truth fiber orientations. Comparison with structure tensor analysis showed that the Frangi filter method better resolves crossing fiber configurations. Being multiscale, the Frangi filter enables the robust determination of the fiber orientation, invariant to the scale of the evaluated WM structure. Also, Frangi filters are specific to bright structures over a darker background, making them ideal for identifying in-plane WM fascicles in S-OCT volumes. Increasing the range and number of scales explored increases the chances of matching tubular structures of corresponding scale. The downside of adding more scales is that it increases computation times. In this work we set the filter scales to {0.5; 1.0; 1.5; 2.0} for real data, as it achieved a good tradeoff between the quality of the reconstruction and execution times. We must however be careful at only allowing scales that would correspond to WM fascicles, to ensure the Frangi filter response is driven by microscale WM structures and not by macroscale structures, e.g. the average orientation of a whole bundle.

#### From orientations to µODF

We used a sliding-window formulation (Zhang et al., 2025) for estimating µODF from the estimated orientations at native resolution, relying on a single parameter *ω* controlling the width of the spatial averaging window. This parameter must be carefully chosen. The simulated dataset consists of mostly straight, macroscale bundles crossing each other at constant angles. As such, a big window extending beyond a complex crossing region and picking up neighbouring coherently-oriented orientations helps regularizing the µODF estimated at the voxel level. However, this loss of specificity to the underlying WM fascicles has a negative impact on tractography reconstructions. This shows that minimizing the angular error locally does not guarantee a good tractography reconstruction globally. Indeed, increasing the spatial averaging window width decreases the mean Dice coefficient and mean overlap. This is because increasing the size of the neighbourhood blurs the estimated µODF, making it harding to precisely follow WM fascicles across complex regions, such as the 3-way splitting between bundles 0, 1 and 2. As such, this parameter should be set to the lowest value allowing for estimating crossing µODF where expected, while remaining as specific as possible to WM fascicles. At the moment, we recommend setting this value empirically based on visual comparison with the underlying data, but future works should focus on developping a metric for quantifying the alignment between the estimated µODF and the underlying fiber configurations.

#### Fiberness measure for anatomical maps definition

On our simulated dataset, we showed that using inclusion and exclusion tracking maps with particle filtering tractography (PFT) to constrain tractography to WM is key for increasing the ratio of valid streamlines. While we built these maps from the ground-truth (GT) bundle masks for the simulated dataset, GT masks are of course not available for real data. Moreover, there are no readily-available methods for reliable brain tissue classification for S-OCT data. In this work, we use the fiberness measure from Frangi filters to estimate the WM probability from S-OCT data. To ensure WM fascicles in low-contrast areas have a similar WM probability than for high-contrast areas, we clipped the fiberness map at the 95^th^ percentile before normalizing to the range [0, 1]. This upper clipping value was chosen empirically based on visual quality assessment of the resulting WM probability map.

#### Particle filtering over deterministic tractography

Our tractography experiments using simulated data showed that PFT with CMC, outperforms local deterministic and probabilistic tractography. Because of the ODF sampling process, probabilistic tractography results in streamlines leaving the boundaries of valid WM fascicles and prematurely terminating outside the WM mask, resulting in many early-terminated streamlines that do not respect the minimum length threshold. With PFT, every stopping event is challenged, with the backtracking mecanism trying to save the streamline by choosing a different high-probability alternative that stays within the tracking mask. For real data, we performed a grid search on the step size, maximum curvature angle and particles count to determine the optimal tracking parameters. Our results showed that using a step size of Δ = 0.010 µm, maximum angle of *θ* = 40^°^ and particles count of 100 maximizes the number of connections between the thalamus and isocortex. This value may be application-dependent, and we recommend performing this parameter sweep when considering other applications, such as for tracking a different structure of interest. An alternative to this parameter sweep could be to use an ensemble tractography approach (Takemura et al., 2016), where tractography outputs obtained from different tracking parameters are combined into a single tractogram. This is a way to increase the number of anatomically valid streamlines, at the cost of an increased number of seeds.

### 5.2. S-OCT tractography: Why?

To the best of our knowledge, we are the first to apply µODF-based tractography to S-OCT acquisitions of the mouse brain. Although they did perform S-OCT tractography, Wang et al. (2015) used a polarization-sensitive system, and applied structure tensor analysis to the retardance signal, which is not measured by our simpler S-OCT system. However, working directly on the S-OCT reflectivity signal has its advantages: the imaging setup is simpler, less expensive and the raw data is less noisy. Compared to targeted microscopy methods such as serial 2-photon excitation fluorescence microscopy or light-sheet imaging, which require tissue clearing and/or viral tracing injections, S-OCT does not require injecting any contrast agent, reducing the sample preparation time from weeks to hours. Furthermore, S-OCT is 3D by nature. As such, the information below the sample blockface can be measured, and isotropic volume reconstruction is possible: in the AMBCA, which uses serial 2-photon excitation fluorescence microscopy, the through-plane resolution is equal to the slicing interval of 100 µm. Also, thanks to its orientation sensitivity, S-OCT enables semi-targeted imaging of the WM. That is, when oriented smartly, S-OCT allows for seeing small WM fascicles hidden inside dense WM bundles, e.g. thalamocortical projections across the CC (recall WM fascicles crossing the CC in Figure 1b). Moreover, as we saw with our comparison against viral tracing experiments from the AMBCA, S-OCT tractography from a single specimen allows for reconstructing thalamocortical projections that would have required the sacrifice of many specimens using targeted microscopy approaches.

While the Frangi filter method is not specific to WM, a proper initialization of tractography and good anatomical maps definitions allow for precisely reconstructing individual WM fascicles connecting the thalamus to the cortex. The biggest promise for microscopy tractography lies in its ability to follow precisely these small WM fascicles invisible at the dMRI resolution. This would help explain how fiber populations are organized at the finest scales, and is key to deriving priors for macroscale tractography algorithms. For instance, if we know where a diffusion MRI tractography streamline comes from as it propagates inside WM, we could imagine having some additional information explaining where it should propagate to based on observations made from S-OCT tractography.

### 5.3. Limitations and future works

#### Simulated Fibercup phantom

One limitation of our simulated Fibercup phantom is that the ratio of crossing fibers to single fibers is only 1%, which is far from what we expect for real S-OCT data. The simulated dataset does not exhibit the complex fiber configurations that we may observe in real S-OCT acquisitions, such as sharp turns and multiscale structures. The simulated dataset does not penalize for the lack of specificity to small WM fascicles within crossing regions as much as it should. Indeed, the crossing configurations can be inferred by examining areas outside those problematic regions. This is why we can always improve the µODF reconstruction metrics by increasing the size of the mean filter. Furthermore, the simulated Fibercup phantom describes complex configurations at the macroscale. However, the problem we are trying to solve is microscale. Another limitation of the phantom is that the simulated S-OCT contrast is not physically accurate, nor does it perfectly mimic the dynamic of a real S-OCT acquisition. While it remains fair for comparing different µODF estimation and tractography methods, there is no guarantee that conclusions drawn from the simulated dataset will transfer to real S-OCT data. Moreover, track-density imaging rasterizes infinitesimally-thin streamline elements over the voxel grid, resulting in 1-voxel wide WM fascicles, which are extremely difficult to follow using a discretized set of propagation directions and discrete step-size. Indeed, previous experiments performed using the original bundle masks as the CMC maps resulted in very poor results (data not shown). This is why we used dilated GT bundle masks to define the CMC maps. However, using dilated WM probability maps results in high bundle overreach, as it allows streamlines to deviate slightly from the true trajectories. As such, the low F1 scores reported may be a consequence of the nature of the simulated dataset rather than a limitation of tractography itself. Instead of using track-density images from the Fibercup phantom, future works should consider designing artificial phantoms better describing the complex fiber configurations observed at the microscale. Track-density images should also be replaced by a model more representative of S-OCT.

#### High rejection rate and false negatives

S-OCT tractography has a high rejection rate of around 98%. Because of noise, it is possible that µODF estimated inside very fine WM fascicles do not always perfectly align with the principal orientation of the structure. Combined with exclusion maps tightly constraining tractography to the structures of interest, taking an invalid step is highly likely. When this happens, backtracking and particle-filtering attempt to save this streamline. However, when the maximum number of trials is reached, the streamline will be rejected before it can connect to a valid endpoint. To reduce the rejection rate, we could try to increase the maximum number of trials for backtracking. For very fine WM fascicles, this may allow to save streamlines and generate less false negatives. Or, we could develop a novel tractography approach where the CMC probability maps are used to weight the local µODF, such that directions leading to the exclusion of a streamline are less probable than valid directions. Another source of error may come from estimating the WM probability maps from the fiberness measure. Here again, the noisy nature of S-OCT may result in holes in the estimated tracking inclusion and exclusion maps, preventing tractography from following WM fascicles along their entire length. To ensure the robust estimation of WM probability, future works should focus on the development of an automated tissue classification methods for S-OCT data.

#### S-OCT artifacts and lack of specificity

The reconstructed S-OCT volume exhibits stripes patterns along the direction of acquisition. This is because the measured signal is better resolved at the surface of an imaged tissue section than below the surface. Artifacts present in the S-OCT volume affect the image derivatives, which are directly used by the Frangi filter method for orientation estimation. Hence, to improve the estimated µODF, effort should be put into improving the S-OCT reconstruction pipeline from Lefebvre et al. (2017). Also, because the variation in signal intensities between the CC and isocortex results in strong image derivatives, Frangi filters estimate µODF running along the CC (anteroposteriorly) instead of perpendicular to it. These orientations translate into streamlines running along the CC and isocortex interface instead of directly projecting to the cortex. This issue may be mitigated by more carefully setting the Frangi filters α, β and γ parameters. For instance, we could perform an exhaustive grid search for setting these values, as in Sorelli et al. (2025). However, in their work, the optimal values are still determined from a qualitative evaluation of the resulting orientations. As such, future works should focus on defining a quantitative metric for assessing the quality of the estimated orientations. Also, to increase specificity to WM, we may consider estimating orientations from an intermediate contrast, estimated from the measured S-OCT reflectivity. This contrast could be, for instance, a WM probability map estimated using a more advanced tissue classification method for S-OCT, as discussed above. We also saw that S-OCT tractography reconstructs out-of-plane WM projections, although the multiscale Frangi filters are only sensitive to bright, in-plane WM structures. In that case, the estimated out-of-plane orientations correspond to the orientations of the surrounding GM rather than that of WM fascicles. Indeed, because out-of-plane WM appears very dark, the variations in intensities between the dark WM surrounding bright GM results in a fiberness greater than 0. However, we saw that this issue rarely occurs.

### Orientation dependency

In this work, we showed that S-OCT tractography enables the reconstruction of anatomically-plausible WM connections when these are aligned with the system imaging plane. However, for WM structures that do not live on a single plane, our current approach for S-OCT tractography is unsuitable. In such cases, we again may need to generate an intermediate WM contrast that would allow for delineating both in-plane and out-of-plane WM fascicles. This could be done by developping a forward model predicting the expected S-OCT intensity for given WM configurations. However, doing so would go against the useful property we used in this work, allowing to disentangle crossing configurations by acquiring data along a carefully chosen orientation. To keep this information intact, future works may instead need to focus on combining acquisitions from different viewpoints (Lefebvre et al., 2024).

#### Reproducibility

We applied S-OCT tractography to a single mouse specimen acquired along the sagittal orientation. While a qualitative comparison suggests that we can reproduce similar tractography results across both brain hemispheres, future work should extend the analysis to more data. We also focused only on thalamocortical projections, as these are mostly aligned with the plane of acquisition, and are relatively short. The same dataset could have been used to reconstruct other anatomical structures, such as the corticospinal tract (CST). However, we anticipate that reconstructing the CST using the proposed approach would be challenging, because it is not as well delineated by GM structures as are the WM projections of the internal capsule, and because it does not connect two GM regions. In that context, updated definitions for the inclusion and exclusion maps for better constraining the tractography algorithm to the CST might be necessary. Applying the proposed methodology for reconstructing other brain structures should also be adressed in future works.

## 6. Conclusion

In this work, we showed that conventional S-OCT enables fiber tractography at the microscale. Our approach combines native-resolution µODF with anatomically constrained particle-filtering tractography, enabling for precisely following WM fascicles and recovering anatomically-plausible pathways between the thalamus and cortex in a whole mouse brain at 10-µm resolution. By enabling label-free reconstruction of fascicles that remain unresolved by diffusion MRI, S-OCT tractography opens a new avenue for linking microscopic fiber organization to macroscale connectivity. Despite many challenges, this work is a first step towards state-of-the-art tractography of microscopy S-OCT data. This work highlights the importance of including anatomical priors for S-OCT tractography, and future work should focus on developping tractography algorithms designed specifically for S-OCT.

## 7. Acknowledgements

The authors would like to thank Frans Irgolitsch, Patrick Lafontaine-Martel, Cong Zhang and Frédéric Lesage for the S-OCT data acquisition and sharing. The authors also thank the Digital Research Alliance of Canada for granting access to their resources for high-performance computing and data storage. Charles Poirier received financial support from the Natural Science and Engineering Research Council of Canada (NSERC) through the Canada Graduate Scholarship – Doctorate program and from Unifying Neuroscience Quebec (UNIQUE) through the Excellence scholarship program. Maxime Descoteaux received financial support from NSERC Discovery Grant (RGPIN-2015-05297) and Institutional USherbrooke Research Chair in Neuroinformatics. Joel Lefebvre received financial support from NSERC Discovery Grant (RGPIN-2020-06109). Laurent Petit received financial support from the French government in the framework of the University of Bordeaux’s France 2030 program/RRI “IMPACT” and from the French National Research Agency (ANR) under Grant ANR-22-CE45-0004 (Project CROSS-TRACTS).

## 8. Declaration of generative AI and AI-assisted technologies in the manuscript preparation process

During the preparation of this work the authors used Large Language Models (LLM) in order to improve sentence clarity and for scientific data visualization. We used Github Copilot Auto model selection, which automatically routes requests to public LLM. In our case, the following models were used: GPT-5.4 mini, GPT-5.3-Codex, Claude Haiku 4.5, GPT-5 mini, GPT-5.6 Luna. LLM were used sparingly and on a case-by-case basis. After using this tool/service, the authors reviewed and edited the content as needed and take full responsibility for the content of the published article.

## A. S-OCT reconstruction

We reconstruct a whole-brain S-OCT volume at 10 µm isotropic resolution, using a Nextflow pipeline (Tommaso et al., 2017) built upon the pipeline from Lefebvre et al. (2017, 2018). The pipeline is divided into two parts: single-slab reconstruction, where the tiles corresponding to a single acquisition depth are assembled into a continuous volume; and whole-brain volume reconstruction, where the reconstructed 3D slabs are stacked on top of each other into a single whole-brain volume.

### A.1. Single-slab reconstruction

Single-slab reconstruction is demonstrated in Figure 15 for an example S-OCT slab from Lefebvre et al. (2024), acquired along the axial orientation. Due to degraded tissue sections, this dataset was not considered for further processing. For each cutting depth, the raw tiles are combined into a mosaic grid (Figure 15, a), using the tile position recorded in the metadata file to localize each tile inside the 3D mosaic. We use the OME-Zarr format specification (Moore et al., 2023), designed by the Open Microscopy Environment (OME) for this purpose. The mosaic grids are then resampled to an isotropic resolution of 10 µm to (i) lower memory requirements for subsequent reconstruction steps and downstream image analysis and (ii) reduce the effect of speckle noise, a common artifact of optical coherence tomography (Varadarajan et al., 2022). As in Lefebvre et al. (2017), we correct for lateral illumination biases using the BaSiC algorithm (Peng et al., 2017) (Figure 15, b) and stitch individual tiles together into a continuous volume using Laplace blending weights (see Figure 15, c.1). The light-beam profile, resulting in additional intensity variations along the depth axis (c.2, red boxes), is compensated. In Lefebvre et al. (2017), the authors fit a shifted Gaussian function on the signal from background voxels to estimate the confocal point spread function (PSF) of the light-beam. However, our experiments showed that this model-driven approach will often diverge when improperly initialized. Furthermore, proper initialization is often challenging when the sample moves slightly out-of-focus because of drifts in the motorized stage responsible for translating the sample under the objective. For this reason, we instead use a model-free approach where the average signal profile in background voxels is used as is to compensate for the light-beam PSF. The effect of correction is shown by the red boxes in (Figure 15, d). We employ a similar approach to compensate for optical attenuation (clearly visible in Figure 15 c.2, d), which consists of rescaling the intensities for each depth such that the maximum foreground intensity is 1 and the median background intensity is 0 (Figure 15e).

**Figure 15.**
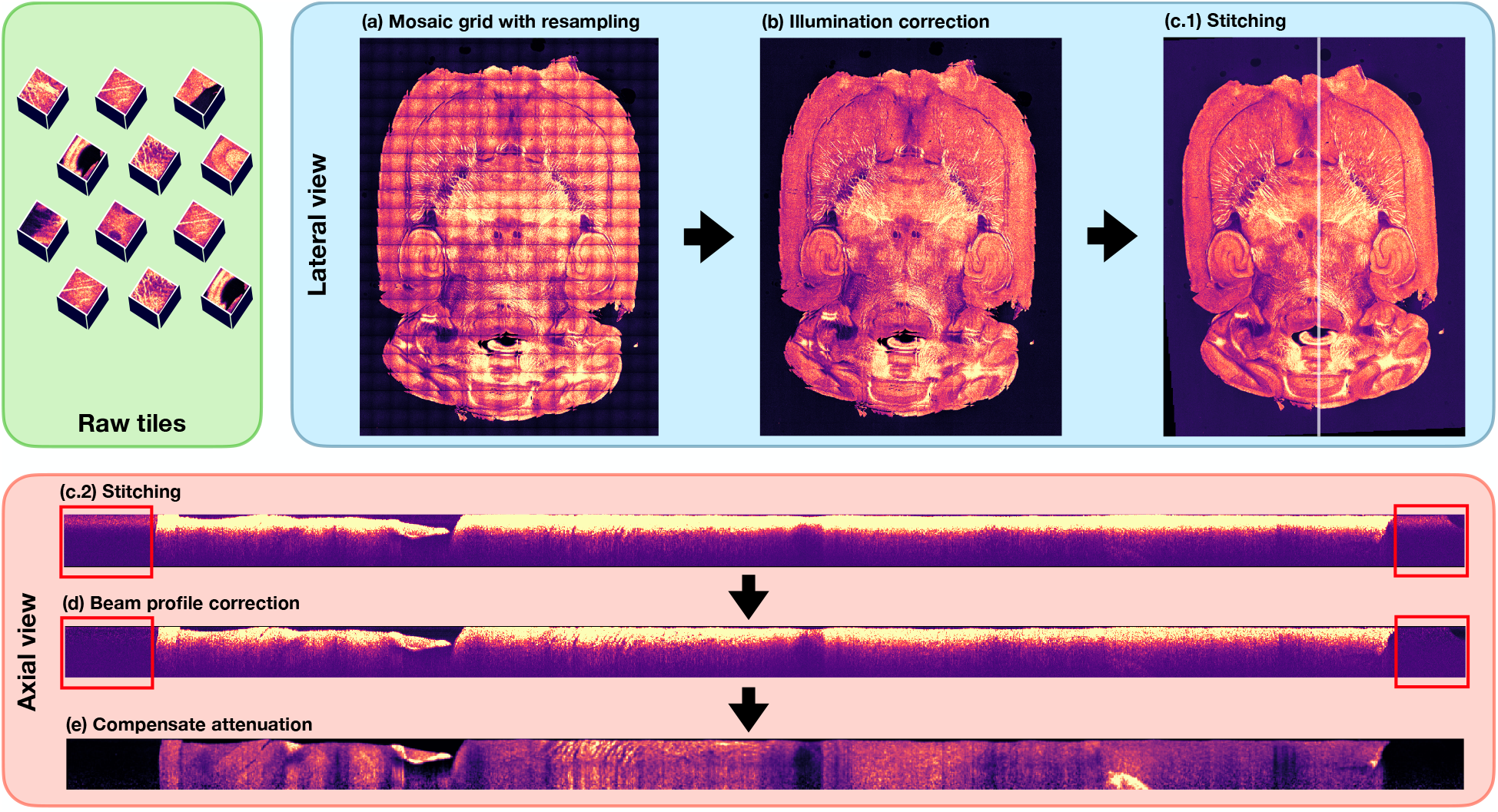
Reconstruction pipeline for an example S-OCT slab (data from Lefebvre et al. (2024)). Green panel: Raw tiles for some acquisition depth. Blue panel: Slab reconstruction in lateral view (in-plane). (a) Raw tiles are assembled into a mosaic grid (OME-Zarr) and resampled to 10µm isotropic resolution. (b) The illumination inhomogeneities inside each tiles is corrected using the BaSiC algorithm. (c.1) The mosaic grid is stitched to remove repeated content at the tiles boundaries. Red panel: Slab reconstruction in axial (side) view for section highlighted in white in (c.1). (c.2) Tiles stiching (same as (c.1)) in side view. (d) The light-beam profile is compensated. The effect of correction is shown by the red boxes in (c.2 and d). (e) The signal attenuation is compensated.

### A.2. Whole-brain reconstruction

Once all slabs are processed, they are assembled into a continous volume covering the whole brain. Because of inaccuracies in the motorized stage translations, simply stacking the slabs at each 200 µm interval is not sufficient for properly aligning the content of each slab. Also, this naive approach underperforms in the presence of tissue deformations occurring over the span of the acquisition. To tackle these issues, we register each slab onto the previous one. Using the recorded motorized stage positions as initialization for our registration algorithm, and the 200 µm slice interval as the initial search depth, we perform 2D affine registration between the top slice of the new slab onto the bottom slice of the previous slab using the Insight Toolkit (ITK) (McCormick et al., 2014). Because of inaccuracies in the motorized stage measurements, we define a tolerance of ±50 µm around the initial search depth of 200 µm. The best registration is the one resulting in the lowest mean-squared error. Then, we stack each slab into a single volume using the estimated transformations. To generate smooth transitions between consecutive slabs, we apply Laplace blending inside an overlap region of 100 µm, as described in (Lefebvre et al., 2017). As a last reconstruction step, we apply the N4 algorithm (Tustison et al., 2010) to the reconstructed volume to correct for variations of intensities between the different tissue slabs. The N4 algorithm models the bias field as a spline defined by control points uniformily covering the volume. Because we expect that the most abrupt variations in intensities happen along the depth axis (stacking axis), we specify a higher number of control points along the depth axis than for in-plane axes. The resulting fully-3D continuous whole-brain S-OCT volume at 10µm isotropic resolution is finally converted to nifti for interoperability with existing tractography softwares.

## B. Structure tensor analysis

Structure tensor analysis (Jähne, 1991) is a gradient-based image analysis method for estimating the principal orientations of an image. Given some image *I* (*x*), we define the structure tensor as

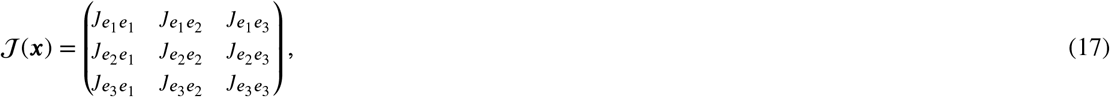

where {*e*_1_, *e*_2_, *e*_3_} *⊂* ℝ^3^ are the standard basis vectors and

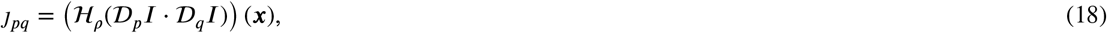

with *D*_*u*_*I* the derivative of *I* along *u* and ℋ_ρ_ a Gaussian window with standard deviation ρ. Let *λ*_1_ ≤ *λ*_2_ ≤ *λ*_3_ be the eigenvalues corresponding to eigenvectors *v*_1_, *v*_2_ and *v*_3_. The vector *v*_3_ associated to the highest eigenvalue *λ*_1_ describes the direction of maximum variation, i.e. the gradient, and *v*_2_ describes the direction of second-to-highest variation, i.e. the tangent to the curve. Then, the vector *v*_1_ corresponding to the lowest eigenvalue *λ*_1_ is the direction of lowest variation of intensities and describes the orienta^1^tion of the structure at *x*.

## C. Tractography scoring metrics (Tractometer)

Here we summarize a few Tractometer metrics which are of interest in this work.

A valid streamline (VS) is a streamline connecting two endpoint regions belonging to a ground-truth bundle (GT). A valid bundle (VB) is the set of all VS connecting two valid endpoints. The bundle-wise overlap (OL) is defined as the number of voxels from the GT mask intersected by a VB, over the number of voxels from the GT mask. The bundle-wise overreach (OR) is the number of VB voxels which do not belong to the GT mask, over the number of voxels intersected by the VB. Finally, the bundle-wise Dice score (F1) is given by

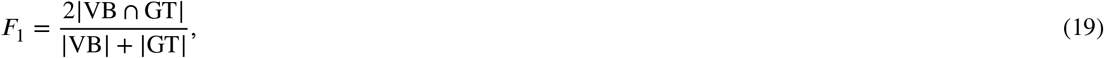

where | ⋅ | denotes the cardinality, i.e. number of voxels, of a set.

## CRediT authorship contribution statement

**Charles Poirier:** Conceptualization, Methodology, Data curation, Software, Writing - Original Draft, Visualization. **Laurent Petit:** Writing - Review and Editing. **Joël Lefebvre:** Writing - Review and Editing, Supervision, Investigation, Resources. **Maxime Descoteaux:** Writing - Review and Editing, Supervision, Resources.

